# Curation of mass spectrometry reference data for improved identification and dereplication of cyanobacterial specialized metabolites

**DOI:** 10.64898/2026.01.12.698359

**Authors:** Franziska Schanbacher, Anne Dax, Michele Stravs, Timo H. J. Niedermeyer, Elisabeth M.-L. Janssen

**Author notes:** co-first authorship (Franziska Schanbacher, Anne Dax).

## Abstract

High-resolution tandem mass spectrometry (HRMS/MS) is a powerful tool for screening organic compounds in complex samples. A critical step in identifying candidate structures is the comparison of sample HRMS/MS spectra with those in reference spectral libraries. The effectiveness of this spectral matching hinges on two key factors: (i) how well the library’s content aligns with the suspect compound list, and (ii) the quality and diversity of the reference spectra for each compound. Yet, the scarcity of natural product reference materials on the market often necessitates non-targeted analysis. In this study, we systematically acquired and curated HRMS/MS reference spectra for specialized metabolites from cyanobacteria, which are vastly underrepresented in current libraries. Previously, MassBank EU included spectra for only 14 such compounds. We have significantly expanded the publicly available data, contributing 2911 unique spectra representing 150 distinct cyanobacterial metabolites. A proof-of-concept analysis demonstrates up to 5-fold increased annotation success and revealed shortcomings in current libraries, underscoring the need for continued data enrichment. In particular, future efforts should prioritize the inclusion of HRMS/MS spectra for diverse adduct ions to improve identification confidence and broaden analytical coverage.

## Introduction

Mass spectrometry (MS), especially high-resolution tandem MS (HRMS/MS), is widely used to screen samples or sample sets for organic compounds, including metabolites, with the example of specialized metabolites (SMs) from cyanobacteria focussed on herein. Moreover, HRMS/MS-based screening approaches have become an integral component of rapid early-stage dereplication workflows, minimizing the risk of rediscovering known compounds in natural product research.^1–3^ Common workflows also incorporate efficient annotation strategies of known compounds to support the comprehensive structure elucidation of yet undescribed congeners.^4^

The two main workflows in mass spectrometry analysis are target analysis and non-target analysis.^5^ Target analysis focuses on compounds for which laboratory methods, including instrument settings, are known and reference standards are available to the analysing laboratory. While reference standards, including isotopically labelled ones, are commercially available for most micropollutants such as pharmaceuticals, pesticides, and industrial chemicals, standards for natural products are scarce and often expensive when derived from purified raw materials rather than chemical synthesis. Consequently, target analysis of natural products is often limited. When reference materials and prior knowledge of optimal instrument methods are unavailable, non-target analysis can be applied. Suspect screening is a type of non-target analysis that searches MS data against a suspect list of compounds whose molecular formulae and chemical structures are known from the literature. Suspect screening enables the researcher to expand beyond the limited number of compounds available for targeted analysis and to increase the likelihood of identifying suspects of interest across a wide chemical space in a given sample. The suspect list has to be defined by the analysing laboratory and can range up to a large and chemically complex lists such as the chemical space provided by PubChem (118 million unique chemical structures, 2025). However, their size and heterogeneity limit screening workflows, so more focused lists are oftentimes preferable. More compact but still broadly applicable resources, such as PubChemLite (566,122 compounds, 2025)^6^, can offer a more manageable alternative, while even more specific focal areas may be selected, for example, bacterial metabolites in The Natural Products Atlas (36‘545 compounds, 2024),^7, 8^ to defined lists based on disciplinary interests, for example, SMs from cyanobacteria in CyanoMetDB (3084 compounds, 2024) as for our example herein.^9, 10^ Even defined lists like CyanoMetDB consist of compounds covering a diverse chemical space and a compromise has to be made regarding the instrument methods selected for the analysis of samples. In suspect screening, while it is not known *a priori* which compounds can be expected in the sample of interest, a suspicion might exist, for example, that the sample contains cyanobacteria and thus some of their SMs are suspected to be present. Hence, the respective suspect list is considered while designing the instrument method, e.g., to define the scan range setting and to include a defined target mass list to trigger fragmentation spectra of those suspects. In contrast to target analysis, the instrument settings will not be optimized for each analyte and rather present a compromise that allows to cover as many compounds from the suspect list as possible, including a selection of conditions for fragmentation (e.g., collision energies). These compromise parameters allow acquisition of HRMS spectra across a wide range of features and are designed to still enable downstream dereplication or tentative annotation, even though the measurement conditions are not individually optimized for each compound.

Following data acquisition, the data analysis workflow of suspect screening comprises broadly two steps. First, the MS features in the sample are compared to the suspect list to identify tentative candidates whose precursor specifications, molecular formula, mass error, and isotope pattern of the MS^1^ scans match. In a second and vital step, a higher confidence of the identity of candidate structures is achieved by matching the measured MS^2^ fragmentation spectra with spectra in a spectral reference library.^11^ No prior knowledge about the fragmentation of the probable structure needs to be known at this stage. Computational tools such as NIST MS search^12^ and The Global Natural Products Social Molecular Networking platform (GNPS, http://gnps.ucsd.edu)^13^ can facilitate automated matching of measured spectra with reference spectra. Further, platforms such as SIRIUS^14–18^ provide both reference spectra search and advanced *in-silico* methods, which include molecular formula annotation, structure database searches for compounds without entries in spectral libraries (i.e., in addition to spectral database searches), and compound class prediction, which can provide additional information in non-target analyses, suspect screening and dereplication approaches.

Spectral matching is a corner stone of successful suspect screening. Its success depends on (i) the compatibility of available compounds in a reference spectral library with the suspect list of interest and (ii) the quality and diversity of the deposited reference spectra for a given compound.^19, 20^ Examples of widely used reference spectra library are MassBank EU, GNPS and the National Institute of Standards and Technology (NIST) reference library. MassBank EU is available since 2011 as one of the first publicly available mass spectral libraries. Since, it developed into a truly open access, open data, open-source resource with highly curated spectra to perform suspect screening.^21, 22^ MassBank is integrated in other resources such as PubChem, the US EPA CompTox Dashboard, NORMAN Database System, and RforMassSpectrometry. As of 2025, Massbank EU offers references spectra for 18‘529 unique compounds and 119‘845 unique spectra that are publicly available.^23^ NIST, which is only commercially available, offers spectra for 51‘501 compounds and 2.4 million spectra.^24^ GNPS contains an open-access knowledge base for community-wide sharing of MS data as reference spectra (GNPS-Collections), including also external data, such as MassBank EU, as well as a crowd-sourced library of community-contributed spectra with a lower level of curation (GNPS-Community). Reference spectra have so far been limited to a subset of compounds: (i) those for which commercially available reference materials exist, and (ii) the few spectra derived from isolated or otherwise annotated compounds, rather than from commercially available standards. The limitation of available reference materials for natural products is a major challenge resulting in the lack of representation in spectral libraries. Before the addition of new spectra outlined herein, cyanobacterial SMs available in MassBank EU comprised only 14 compounds, including eight microcystin congeners (microcystin-LR, -LA, -LF, -LY, -LW, -RR, -YR, [dAsp^3^, Dhb^7^,]-microcystin-RR) as well as nodularin-R, aerucyclamide A, oscillamide Y, and anabaenopeptins A, B, and NZ857. These entries comprise a total of 215 unique spectra, for [M+H]^+^ and [M-H]^-^ precursor ions with a range of collision energies (15-180%) and MS^2^ resolution ranging from 7‘500 to 35‘000 (Table S1).

Here, we systematically acquired and processed reference spectra of cyanobacterial SMs, not only from reference materials but mainly from semi-purified or crude biomass extracts, to enable suspect screening via spectral library matching. We followed established procedures for measurement and data processing and included compounds amenable to reversed-phase liquid chromatography. Detection was performed using HRMS/MS with electrospray ionisation (ESI Orbitrap). Instrument settings covered typical non-target screening conditions, including positive and negative ionisation modes and a wide range of collision energies. Raw data were processed with the open-source R package RMassBank (https://github.com/meowcat/RMassBank-scripts) for automated recalibration, clean-up, intensity correction, metadata retrieval, and MassBank export.^25^ In total, 2911 unique spectra of 150 SMs were uploaded to MassBank EU. A proof-of-concept study identifies missing compound families or representatives.

## Materials and Methods

The samples were provided either as reference materials or biomass extract containing known SMs from cyanobacteria based on previous NMR or MS analyses by collaborators of CyanoMetDB (see acknkowledgments). The teams provided metadata for each sample including the cyanobacterial species as the producing organism, structural information of the metabolites (CyanoMetDB_ID, molecular formula, SMILES), and the expected precursor ion (adduct, charge state, m/z). Materials were stored at -20°C until analysis.

### HPLC-HRMS/MS analysis

For 336 compounds, HRMS data acquisition was performed on an Orbitrap Exploris 240 mass spectrometer equipped with a heated ESI interface coupled to a Dionex UltiMate 3000 RS pump HPLC system (Thermo Fischer Scientific). The following chromatographic parameters were used: XBridge C18 column (50 × 2.1 mm, 3.5 µm, 100 Å, Phenomenex), binary gradient from 10 to 50% in 4 min, and from 50 to 95% for 13 min MeOH in H_2_O (0.1% formic acid each) at 0.2 mL/min, 95% MeOH in H_2_O for 8 min. HRMS data acquisition: pos. and neg. ionization mode, ESI spray voltage 3.5 kV and -2.5 kV, capillary temperature 320 °C, tube lens 70 V, sheath gas flow rate 40 L/min, auxiliary gas flow rate 10 L/min. Full scan spectra were acquired from *m/z* 100 to 1000 with a resolution of 120‘000 at *m/z* 200. MS/MS spectra were acquired in data-dependent acquisition mode (dd-MS^2^), a range of 9 MS/MS experiments in each pos. and neg. mode with a HCD energy of 15%, 20%, 25%, 30%, 40%, 50%, 60%, 70%, 80% at a resolution of 15‘000 at compound-specific varying *m/z* (100-1000). Isolation window 1.0 Da, automatic gain control (AGC) target 5 × 10^4^. All measurements were recorded in profile mode with two consecutive scan range settings: First, with an automatic scan range, starting at the precuror *m/z* and scanning down to 1/15 of the precursor *m/z*, and second, fixing the starting *m/z* to 40, without considering the precursor *m/z*. In total, for each compound, 36 unique spectra types were considered with 9 collision energies, 2 ionization modes (pos. and neg.), and 2 MS/MS scan modes (auto and first fix mass *m/z* 40) (Figure 1).

**Figure 1.**
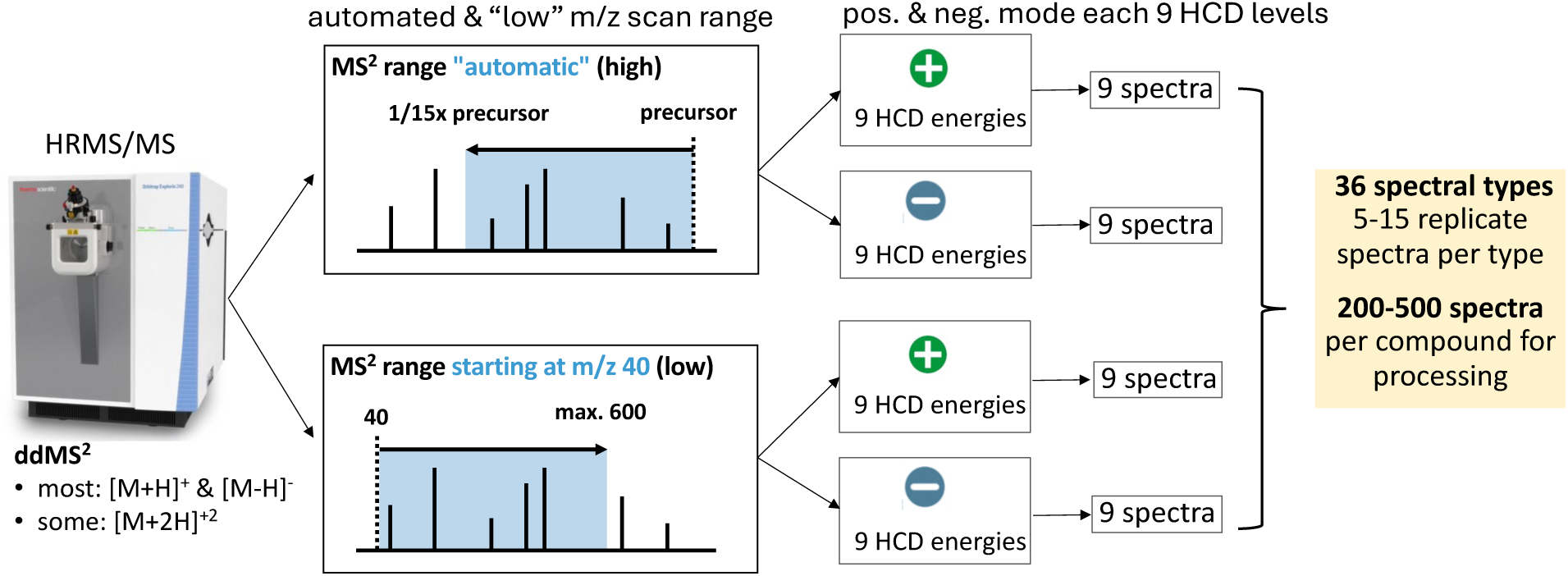
Illustration of MS instrument settings including 2 scan range modes with automatic settings adjusted for each precursor or fixed first mass at *m/z* 40, positive and negative ionization mode with 9 collision energies in each mode (15, 20, 25, 30, 40, 50, 60, 70, 80%) to generate up to 36 unique spectral types with 5-15 relicates per type and a total of 200-500 spectra per compound for data processing.

In addition, 54 compounds were analyzed using an Acquity UPLC HSS T3 analytical column (100 × 3 mm, 1.8 µm, Waters) to improve chromatographic separation. The following chromatographic parameters were used: binary gradient from 15 to 100% MeOH in H₂O (0.1% formic acid each) over 25 min at 0.2 mL/min, followed by 98% MeOH in H₂O for 6 min at 0.65 mL/min. Compared to the method described above, collision energies of 15–70% were used, with an additional 35% step. Otherwise, identical HPLC and HRMS/MS settings on the Orbitrap Exploris 240 were used as described above. In addition to [M+H]^+^ and [M–H]^-^, [M+2H]^2+^ precursors were considered increase chances for adequate HRMS^2^ spectra.

Additional 75 compounds were analysed with slighty different setting on an QExactive Plus mass spectrometer equipped with a heated ESI interface coupled to an UltiMate 3000 HPLC system. The following chromatographic parameters were used: Kinetex C18 column (50 × 2.1 mm, 2.6 μm, 100 Å, Phenomenex), a binary gradient equivalent to the previously described method, with MeCN replacing MeOH, at 0.4 mL/min. HRMS data acquisition: pos. and neg. ionization mode, ESI spray voltage 3.5 kV and −2.5 kV, capillary temperature 350 °C, sheath gas flow rate 50 L/min, auxiliary gas flow rate 12.5 L/min. Full scan spectra were acquired from *m/z* 100-1250 with a resolution of 70‘000 at *m/z* 200, AGC target 1 ×10^6^, maximal injection time 50 ms. MS/MS spectra were acquired in dd-MS^2^ mode, HCD range of as for the other measurements from 15% to 80% at a resolution of 17‘500 at *m/z* 200, an AGC of 2 × 10^5^, and a maximal injection time of 50 ms.

### Data processing

The raw data files were converted to mzML with ProteoWizard MSConvert (3.0.22116-18c918b) using vendor centroiding. Metadata of the analysed materials and compounds was prepared in a CSV table containing the CyanoMetDB compound ID, the compound name, SMILES code, molecular formula, and retention time. A settings file was prepared defining the processing options and generic metadata to be added to the database spectra. The raw data files and experimental settings file were used as input for a data processing workflow based on RMassBank^25^ with extensions to further improve data quality and simplify manual curation (https://github.com/meowcat/RMassBank-scripts/).

Briefly, the data processing workflow consisted of the following steps: (1) Extraction of MS/MS spectra based on the compounds specified in the metadata. (2) Assignment of molecular formula to each fragment with large tolerance, using a ±15 ppm window and a ±10 ppm window for low mass range (<= m/z 120) and high mass range. (3) Calculation of recalibration curve of mass error against m/z values with uniquely assigned fragments, using peaks with intensities >=10^3^. (4) Recalibration of all spectra with calibration curve. (5) Subformula re-assignment with smaller tolerance (less than 5 ppm deviation). (6) Unassigned fragments were checked allowing for N2 and/or O addition to detect fragment adducts with collision gas. (7) Fragments occurring in at least two spectra were retained. (8) To ensure the accuracy of the extracted spectra, in particular to eliminate spurious formula matches of low-mass fragments, an orthogonal analysis based on EIC correlation was performed. For all precursor and fragment ions, EICs were extracted, and the correlation/dot product of fragment to precursor ion EIC was calculated. Results from formula assignment were used to automatically determine a cutoff for EIC correlation: the cutoff was chosen such as to maximize the F_1.5_ score for predicting formula match from EIC correlation. (9) Spectra were reviewed in a graphical user interface (Figure 2) which allowed to review formula match and EIC correlation for individual fragment of each spectrum, as well as to include or exclude spectra or entire compounds, or adjust correlation thresholds to a custom value per spectrum or compound if required.A quality threshold was applied based on the EIC correlation of fragments and the precursor. In the spectral plot, all fragments with poor EIC match (below the quality cutoff) were plotted in red in case they were assigned a molecular formula, or black if no molecular formula was assigned. Both types were removed and thus not part of the spectrum in the final export. Fragments illustrated in green matched with precursor EIC, but have no molecular formula assigned, which can occur when a second precursor is in the isolation window, or at the bottom of the mass range outside the well-calibrated domain. These fragments were included, but called for review of the purity of the MS^2^ spectrum. For standard reference materials that could be measured with an approriately high concentration and purity, a stricter EIC match threshold could be applied. When working with compounds at a large range of intensities from biomass extracts or semi-purified extract fractions, individual attention was given to each compound to make a case by case decision on the threshold for eliminating fragments.

**Figure 2.**
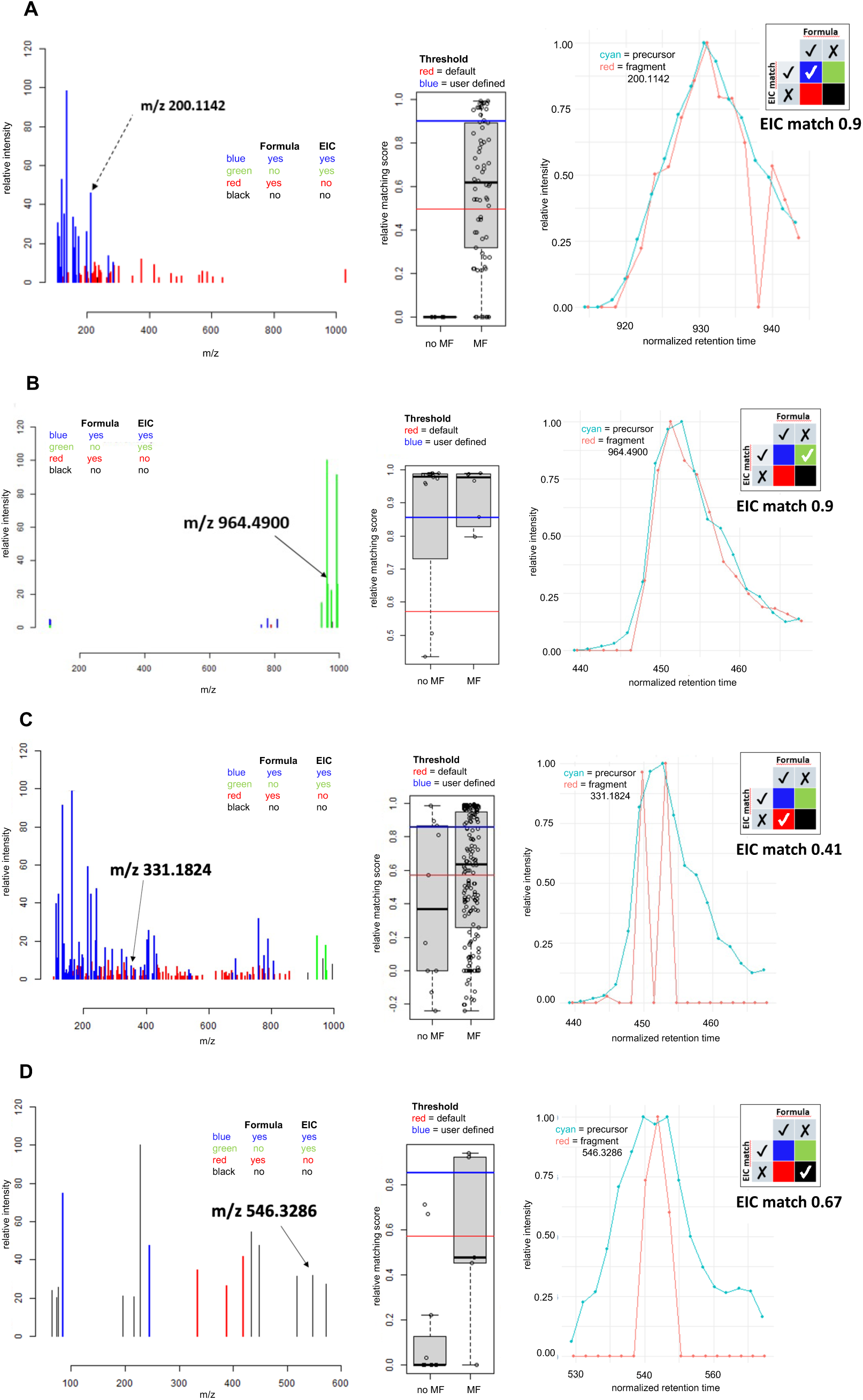
Examples of data represented in the viewer graphical user interface comparing the correlation of fragment to precursor ion EIC. All fragments in the mass spectrum (left) are categorised with **(A)** molecular formula assigned and with good EIC (blue), with **(B)** no molecular formula assigned and good EIC match (green), with **(C)** molecular formula assigned and poor EIC match (red), and with **(D)** no molecular formula assigned and poor EIC match (black). The range of goodness of the relative EIC match for all fragments with and without molecular formula (MF) assinged are shown in a box plot (middle) with the quality cutoff for good EIC match in red being the viewer default and blue being a user defined threshhold selected for each compound. The precursor EIC (cyan) can be compared with each fragment EIC (red) as relative intensity against retention time (right).

Finally, all selected spectra were exported to MassBank records, combining processed spectra data with experiment metadata from the settings file (e.g., MS type MS^2^, chromatographic details), and compound metadata retrieved from CyanoMetDB and internet services. Each record contains the team (author) and their prior evaluation of the confidence level of the reported compound.

Level 1 was used when the compound in the supplied material for analysis was at a reference material purity (>95%), and previous MS^2^ annotation and NMR studies confirmed the identity. Level 2a was used when the compound‘s spectrum could be match with an available refereence spectrum, and Level 2b was used when the compound was previously identified in the same type of material as the one provided, e.g., extract of the same cyanobacterial strain. The identification may have been done by manual annotation of MS^2^ spectra recorded by the collaborating laboratories. When a peer-reviewed reference was available presenting the annotation, a citation was also included in the respective MassBank record. Level 3 was used when the compound identity was inconclusive regarding other structural isomers of the same compound class from previous HRMS^2^ annotations of the same sample material. In these cases, the same MassBank record was associated with all isomer names. For all records derived from non-reference materials, the spectra were marked with “TENTATIVE” in the record title, in line with other existing non-reference records in MassBank.

### HPLC-HRMS/MS Data Acquisition for proof of concept and case study

HRMS data analysis was carried out on 15 cyanobacterial biomass extracts selected based on a priori knowledge of their SMs, with reference compounds included to evaluate annotation and recovery rate for a proof-of-concept study (**A**) and on 4 additional cyanobacterial biomass extracts selected for a case study (**B**). The dataset comprised extracts from (A) *Fischerella*s sp. (ambigols,^26^ tjipanazoles^27^), *Tolypothrix* sp. (cyanobacterin and analogues), *Nostoc* sp. (nostotrebin 6 and congeners^28^, cryptophycins), *Hapalosiphon* sp. (hapalindoles^29^), *Limnothrix* sp. (acutiphycin), *Microcystis* sp. (aerucyclamides, microginins, microcystins^30, 31^), *Planktothrix* sp. (anabaenopeptins), *Cylindrospermum* sp. (cylindrofridins), and *Scytonema* sp. (Scyptolin A and B) (Table S2), and (B) from *Microcystis* sp. (PCC7806: microcystins, cyanopeptolins, cyclamides), *Planktothrix* sp. (K-0576: microcystins, anabaenopeptins) and *Dolichospermum* sp. (NIVA-CYA 269/6: microcystins, cyanopeptolins, cyclamides) (Table S3)^32^. The HRMS data acquisition was performed either on a (**A**) Q Exactive Plus mass spectrometer or (**B**) Orbitrap Exploris 240 mass spectrometer, both equipped with a heated ESI interface coupled to an UltiMate 3000 HPLC system. The following chromatographic parameters were used: Kinetex C18 column (50 × 2.1 mm, 2.6 µm, 100 Å, Phenomenex), (**A**) binary gradient from 5 to 100% MeCN in H_2_O (0.1% formic acid each) at 0.4 mL/min in 16 min, 100% MeCN for 4 min; (**B**) binary gradient 7% for 3 min, from 7 to 30% in 4 min, and from 30 to 100% in 23 min MeOH in H_2_O (0.1% formic acid each) at 0.255 mL/min, 100% MeOH for 7 min. HRMS data acquisition: (**A**) pos. and neg. ionization mode, ESI spray voltage 3.5 kV and -2.5 kV, capillary temperature 350 °C, sheath gas flow rate 50 L/min, auxiliary gas flow rate 12.5 L/min, (**B**) pos. ionization mode, ESI spray voltage 3.5 kV, capillary temperature 320 °C, sheath gas flow rate 40 L/min, auxiliary gas flow rate 10 L/min. Full scan spectra were acquired from: (**A**) *m/z* 133.4 to 2000 with a resolution of 35’000 at *m/z* 200, automated gain control (AGC) 5 × 10^5^, maximal injection time 120 ms, (**B**) *m/z* 110 to 1500 with a resolution of 120‘000 at *m/z* 200, AGC of 2.5 × 10^5^, maximal injection time 50 ms. MS/MS spectra were acquired (**A**) in data-dependent acquisition mode (dd-MS^2^), stepped collision energy of 30, 60, and 75 eV (resulting at 55 eV), a resolution of 17500 at *m/z* 200, an AGC of 2 × 10^5^, and a maximal injection time of 75 ms, (**B**) in data-dependent acquisition mode (dd-MS^2^), CyanoMetDB as the targeted mass list, 3 MS/MS experiments with HCD 15%, 30%, and 45%, a resolution of 15‘000 at *m/z* 200, an AGC set to standard (set in automated fashion dependent on scan type), and a maximal injection time set to auto (system sets the maximum injection time available according to the transition length, optimizing between sensitivity and scan speed). A TopN experiment (N = 5, loop count 5) was implemented for triggering the dd-MS^2^ acquisition for both (A and B).

### Data processing proof of concept data

#### File Conversion

Raw mass spectrometry data files were converted from .RAW to .mzML format using MSConvert from ProteoWizard (version 3.0).^33^ A scan polarity filter was used during data conversion to separate positive ion mode scans from negative ion mode scans, facilitating a more targeted analysis in the subsequent data analysis steps.

#### Feature-Based Molecular Networking (FBMN) and SIRIUS Analysis

To facilitate FBMN analysis, the converted MS data were processed using MZmine (versions 3 and 4.2) with a workflow designed and executed through the MZWizard tool to automate the feature extraction and alignment.^34^ For mass detection, the following parameters were used to discard low-intensity signals: noise level thresholds 6.00 (MS) and 0.00 (MS^2^) . Chromatogram building was performed with the following parameters: minimum consecutive scans 4, minimum absolute height 1.0E6, *m/z* tolerance 10 ppm.^35^ Chromatographic peaks were smoothed using a Savitzky-Golay filter (window size 5 points) to reduce noise and refine the peak shape. The Join Aligner module for peak alignment was used with the following parameters: retention time tolerance 0.4 min, *m/z* tolerance 5 ppm. MGF files were exported from MZmine 4.2 for analysis in SIRIUS. Molecular formula determination, molecular structure database searches, and spectral library matching were conducted using SIRIUS 6.0.1 and 6.3.2 (CSI:FingerID).^14–16^ For the structure database search, a custom suspect database was compiled from the SMILES strings provided by the current version of CyanoMetDB (version 3.0).^9, 36^ Spectral library matching was restricted to the following two custom libraries of reference spectra: (i) spectra from the MassBank database (accessed March 2025), (ii) spectra acquired and recorded within this study. The following settings were applied in the compute dialog. Global configurations included: Orbitrap as instrument with an HRMS^2^ mass accuracy of 5 ppm, Fallback Adducts set to [M+H]^+^, [M+Na]^+^, [M+K]^+^, and [M-H_2_O+H]^+^ for pos. mode, and [M-H]⁻ and [M+CH_2_O_2_-H]⁻ for neg. mode, Search DBs included the custom/suspect databases and the default SIRIUS databases PubChem, CHEBI, COCONUT, GNPS, HSDB, LOTUS, Maconda, SuperNatural, and TeroMOL. Spectral library searching was performed using default settings. Molecular formula identification was conducted using the de novo + bottom-up strategy, with the element filter set to default values and Br and Cl additionally allowed, and autodetection of Mg, B, Fe, Zn, and Se disabled. The analysis results were exported, and summaries of the structure database searches and spectral library matching were incorporated into the FBMN analysis during visualization in Cytoscape. All annotations were manually reviewed prior to network visualization to ensure their plausibility and consistency. The molecular networks for FBMN analysis were generated using GNPS (http://gnps.ucsd.edu).^13^ All HRMS² fragment ions within a 17 Da window of the precursor *m/z* were excluded from the data set. HRMS^2^ spectra were further filtered to select the six most prominent fragment ions within a 50 Da window. Precursor ion mass tolerance was set to 0.02 Da, and the same tolerance was applied to HRMS^2^ fragment ions. Networks were constructed by retaining edges with a cosine score > 0.6 and a minimum of four matched peaks. Edges were retained only if both nodes appeared in each other’s top ten most similar nodes list. The maximum size of any molecular family was limited to 100, and the lowest scoring edges were removed until each molecular family was below this threshold.^13, 37^ The resulting molecular networks were visualized and analyzed using Cytoscape (version 3.10.2).^38, 39^

## Results and Discussion

### Acquisition and Processing of Reference Spectra

In total, 411 unique compounds were analysed. Depending on the chromatigraphic peak width, 200-500 individual spectra were recorded per compound and spread over 36 spectra types (9 collision energies, 2 ionization modes, and 2 HRMS^2^ scan modes, (Figure 1)) with 5-15 replicates for each spectrum type. Of all analyzed samples, 365 compounds were well detected, 277 compounds passed the data processing, and 150 compounds passed a final manual check by the expert who supplied the material and was familiar with the respective expected tandem mass spectrum (Figure 3A). A total of 2911 unique new spectra of these 150 compounds could be uploaded to MassBank EU. An overview is provided in Table S4. The main reasons why compounds were rejected during data processing were insufficient intensity of the precursor peak and/or too many interferences from the background. Despite many spectra not passing quality control, we significantly increased the coverage of SMs from cyanobacteria in MassBank EU, notably expanding the chemical space. We provided new spectra for 11 compounds that were previousely represented in MassBank EU (marked with # in Table S4), and introduced 140 compounds as novel entries into the spectral library. For example, 21 microcystin congeners were added to the existing eight. Many new compound classes are represented for the first time, including cyanopeptolins, microginins, aeruginosins, or spumigins, among many others. Overall, we increased the number of unique spectra of SMs from cyanobacteria from 215 to 2911 with higher resolution and wider range of collision energy range.

**Figure 3.**
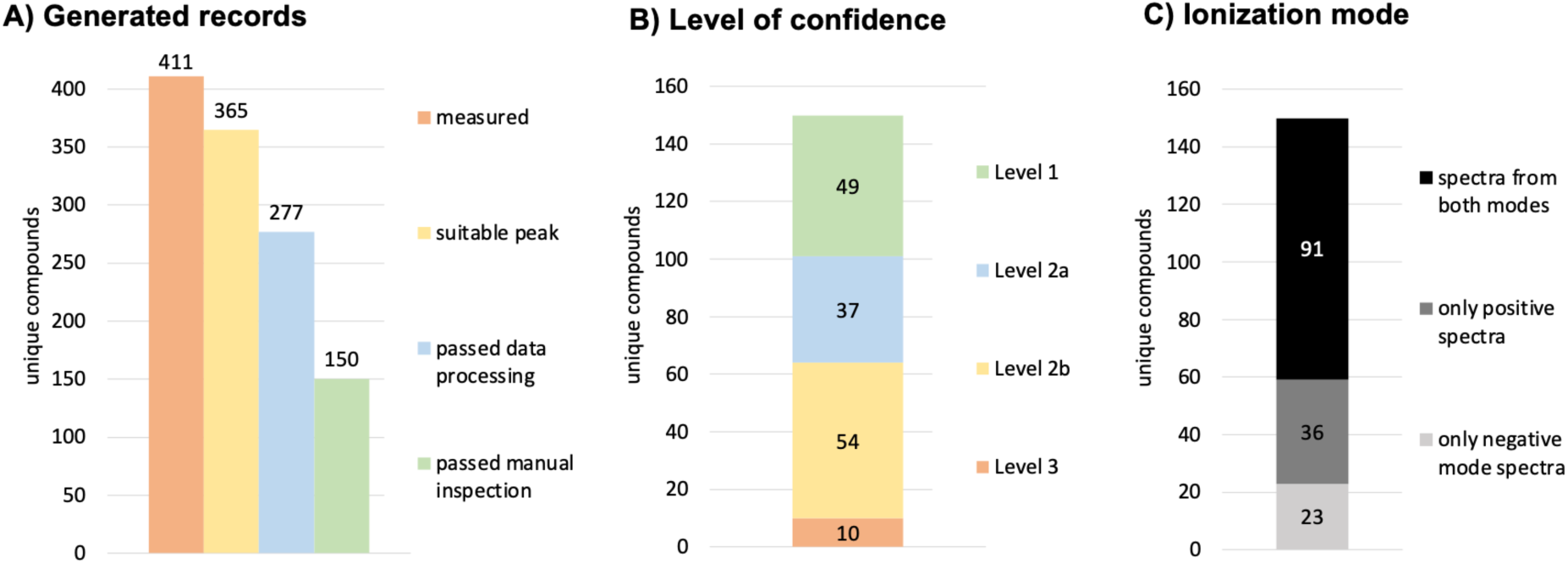
Overview of results. (A) 411 unique compounds were analyzed, 365 compounds were well detected, 277 compounds passed data processing, 150 compounds passed manuel inspection and were approved for record generation and upload to Massbank EU. (B) Level of confidence reached for approved records range from 49 compounds on Level 1, 37/57 compounds of Level 2a/2b, and 10 compounds on Level 3. (C) Availability of ionization mode in spectra types showed both modes for 91 compounds, only positive mode for 36 compounds, and only negative mode for 23 compounds.

Of the 150 unique compounds, 49 compounds were classified at the confidence level 1 (reference materials), 37 compounds at the confidence level 2a, 54 compounds at the confidence level 2b and 10 compounds at the confidence level 3 (Figure 3B). The confidence level is marked in the MassBank EU record for each compound. All records derived from non-reference materials (lower than Level 1) are marked with “TENTATIVE” in the record title which has to be taken into account when library matches are achieved during data processing. Care should be taken with confidence Level 3 library matches, since the annotation of the compound identity was partially inconclusive regarding other structural isomers, and the same MassBank EU record was associated with all isomer names in the database (see last 10 entries in Table S4). Note that these records are not counted multiple times when referring to unique compounds herein. Regarding the ionization mode, 23 compounds could only be detected in negative mode, 36 could only be detected in positive mode, and 91 compounds were detected in both ionization modes (Figure 3C).

### Proof-of-Concept Study

In a proof-of-concept study, the impact of the newly deposited HRMS^2^ reference spectra on the annotation of compounds known by the authors to be present in specific cyanobacterial extracts was investigated (Table S2). Annotation rates were compared before and after adding the reference data discussed above, demonstrating how the expanded spectral library influences the identification of cyanobacterial SMs. Recovery rates were assessed by determining whether these newly added reference compounds, several of which are present in the analyzed extracts (Table S2), could be rediscovered and, if so, with what degree of spectral similarity between the reference spectra and the measured HRMS^2^ spectra. In total, FBMN of the positive-mode data of the selected strains revealed 3351 nodes, of which 2233 features were organized into clusters. Of these, 26 nodes were annotated via spectral-library matching, while 139 were annotated through structure-database searches. Notably, 16 of the 26 spectral-library-matching annotations - and a total of 165 annotations (structure-database searches + spectral-library matching) - could be achieved exclusively through the newly recorded reference spectra generated in this study (Table S5). The remaining 10 annotations were based on spectra already present in the previous MassBank dataset. Of these, six were annotated exclusively through MassBank data via spectral-library matching in SIRIUS, whereas the other four could likewise be annotated using the corresponding user-contributed cyanobacterial spectra available on the GNPS platform. Moreover, one annotation (Microginin FR1, Figure 3F) obtained through the structure-database search was confirmed using a user-contributed spectrum from the GNPS platform. Furthermore, the newly added reference data enabled the identification of five compound-family clusters that had previously remained undetected (Figure 4, D-G). Using the annotated SMs as key nodes in the FBMN analysis, the confidence of structure database-based annotations for corresponding congeners could be improved (Figure 4, B-G). The inclusion of the new reference spectra led to a substantial increase in the overall annotation rate, from roughly 0.3% to 0.8%, a 2.5-fold increase. When cluster-based annotation relationships are considered - such as in the cryptophycin cluster (Figure 4E) - the total annotation coverage across the dataset increases to approximately 1.6%. Seven additional annotations based on the newly recorded reference spectra were enabled through the FBMN analysis of the negative-mode data (Figure 5).

**Figure 4.**
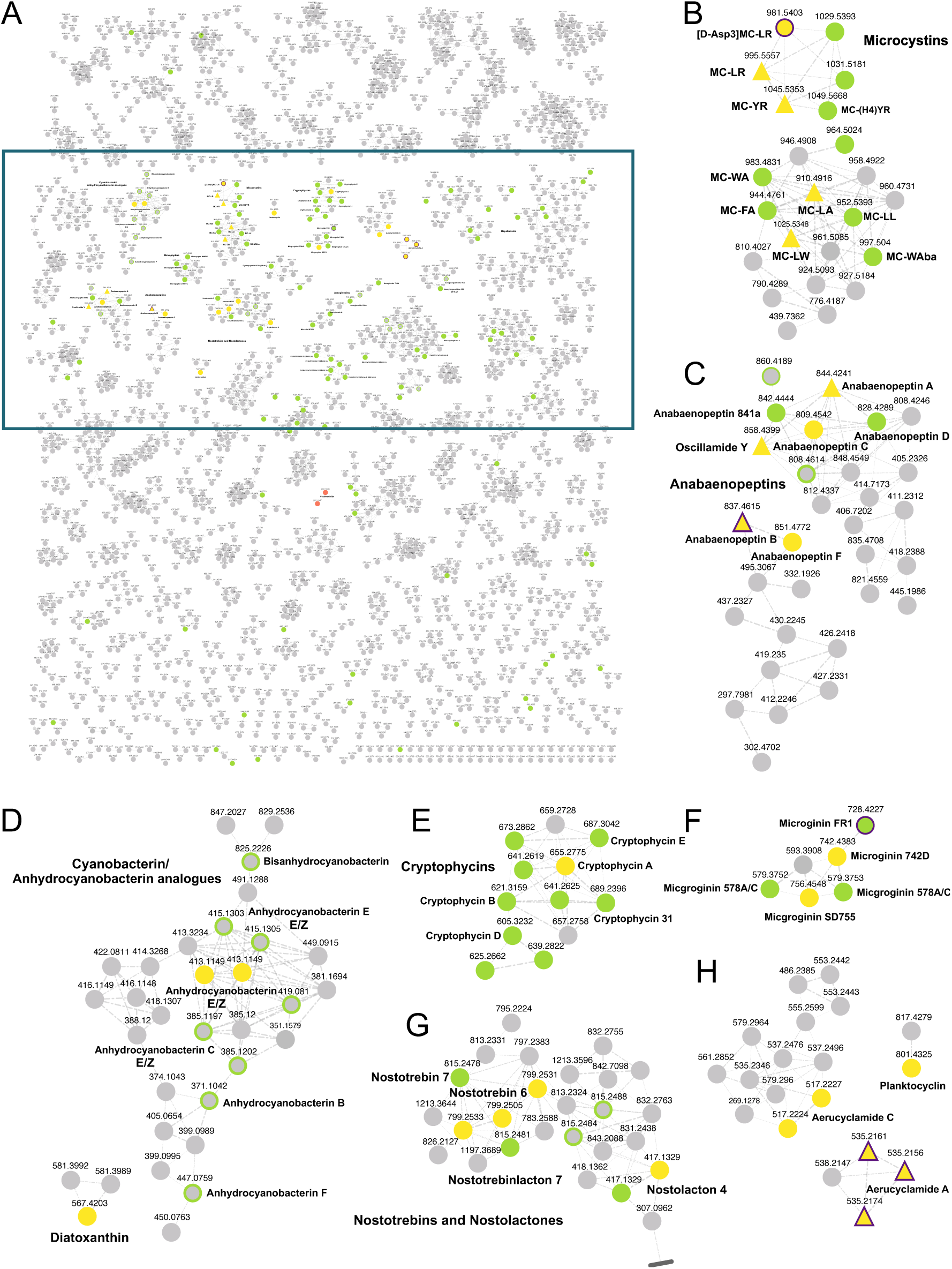
**A** Feature Based Molecular Networking (FBMN) analysis (overview) of a sample set of cyanobacterial extracts with major compounds known a priori to the authors (Table S2) for chemical space visualization, positive mode data. Compound annotation was achieved through spectral library matching and structure database search using the SIRIUS platform. Nodes connected for cosine scores ≥0.7, edge width correlating to cosine scores. Area of clusters with features annotated via spectral library matching is highlighted. Annotations obtained through spectral library matching are highlighted in yellow, and those derived from structure database searches in green. Triangular nodes indicate spectra previously included in the MassBank database, while nodes outlined in purple represent features additionally annotated via GNPS. Features without annotation, either corresponding to cyanobacterial SMs or uncharacterized compounds in general, are shown in gray. To enhance readability, single unconnected nodes were excluded from the FBMN network representation. Cluster of **B** microcystins, **C** anabaenopeptins, **D** cyanobacterins^40^ **E** cryptophycins, **F** microginins, **G** nostotrebins/nostolactons (the cluster is not shown in its entirety due to space constraints; no annotations were associated with the omitted features), and **H** aeruclycamides, where spectral annotation of main compounds was enabled by newly acquired HRMS^2^ spectra recorded in this study, demonstrating enhanced annotation coverage.

**Figure 5.**
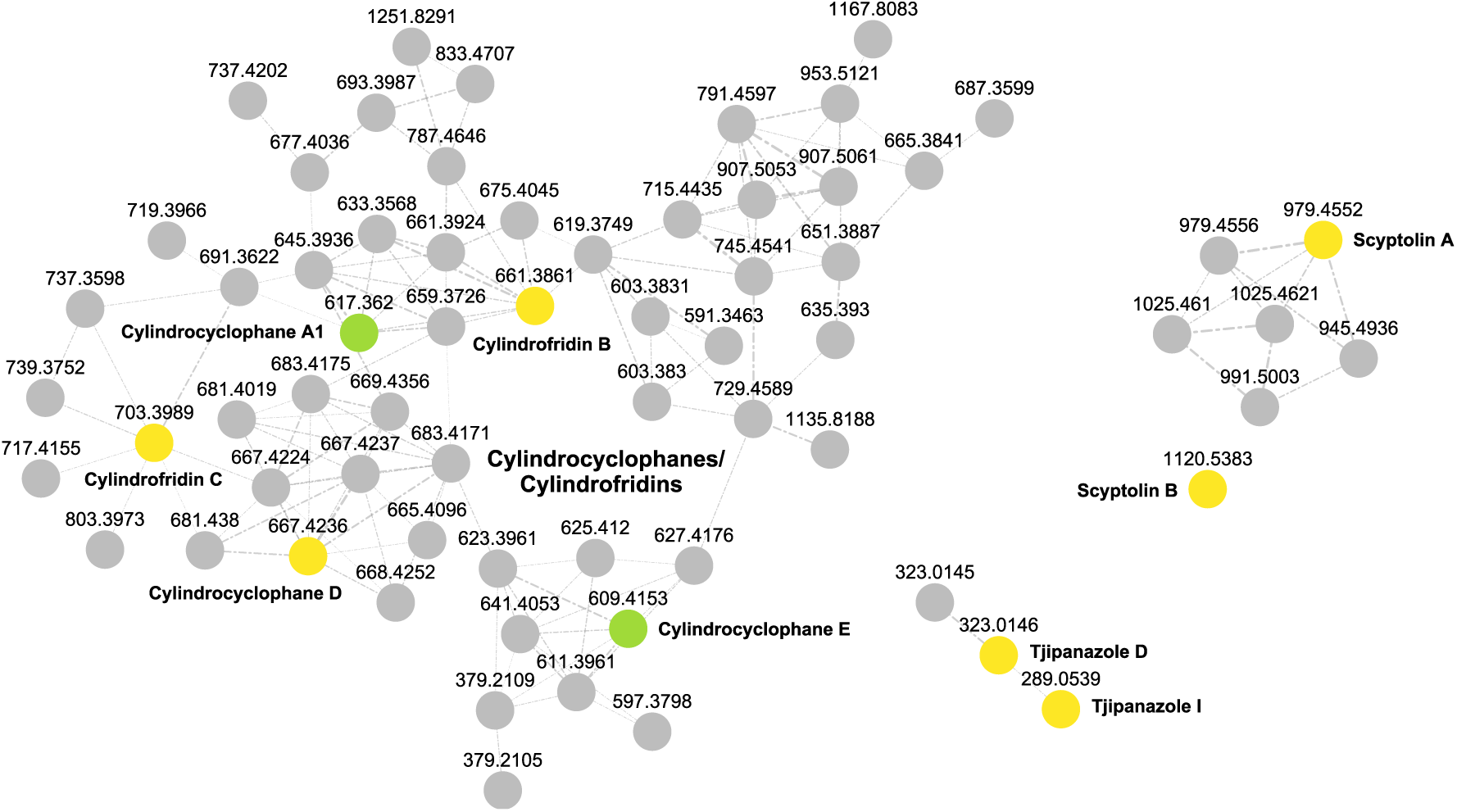
Feature Based Molecular Networking (FBMN) analysis (selected clusters) of a sample set of cyanobacterial extracts with major compounds known a priori to the authors (Table S2), negative ionization mode data. Compound annotation was achieved through spectral library matching and structure database search using the SIRIUS platform. The clusters corresponding to cylindrocyclophanes/cylindrofridins, scyptolins, and tjipanazoles were identified exclusively in negative ionization mode data. Nodes connected for cosine scores ≥0.7, edge width correlating to cosine scores. Annotations obtained through spectral library matching are highlighted in yellow, and those derived from structure database searches in green. Features without annotation, either corresponding to cyanobacterial SMs or uncharacterized compounds in general, are shown in gray.

Except for one compound, all expected SMs were successfully rediscovered (Table S5). Although reference spectra for Acutiphycin were acquired, no corresponding features were identified in the extracts. Pure substance was used for reference spectrum acquisition, and inclusion lists were generated based on [M+H]⁺ or [M–H]⁻ ions. However, Acutiphycin tends to form adducts and undergo water loss during ionization. In some cases, the [M+H]⁺ signal was very low, resulting in insufficient ion intensity for MS^2^ fragmentation under the applied method. Furthermore, the HRMS^2^ spectra differ between the [M+H]⁺, [M+Na]⁺, and [M-H_2_O]⁺ ions. This example highlights the necessity of accounting for potential adduct formation when acquiring reference spectra in future studies.

The analysis also revealed remaining gaps in the reference data, pinpointing which representative members of each compound family should be prioritized for future measurements. For example, the microginin cluster highlights the need for additional reference data. In untargeted screening, acquisition parameters always involve compromises and are not optimized for specific compound classes. Except for Microginin SD755 and Microginin 742D, no other microginin congeners could be annotated from spectral data (Figure 4F), and their matching scores are low (Table S5, 32 % and 24%). In addition to the two annotated microginins, other microginins are expected to be present. Although spectral annotation for these additional compounds was not possible, the low mass deviations between exact and accurate masses, together with their co-clustering in the FBMN with the annotated microginins, suggest that these features likely correspond to the microginins of interest. These findings not only highlight the necessity of optimizing acquisition parameters to reliably detect and annotate microginins, but also the importance of systematically expanding reference data. As illustrated in Figure 6, the lack of reference spectra affects not only microginins but also other relevant cyanobacterial peptides.^41^ The clusters shown in that figure correspond to members of several compound families for which spectral references are still unavailable, highlighting the demand for additional data to enable more confident annotation across diverse compound classes, e.g., the large compound family of hapalindoles, fischerindoles, and welwitindolinones^29, 42–46^ as well as micropeptins,^47^ cyanopeptolins^48^ (see also Figure 7D), or aeruginosins.^49^

**Figure 6.**
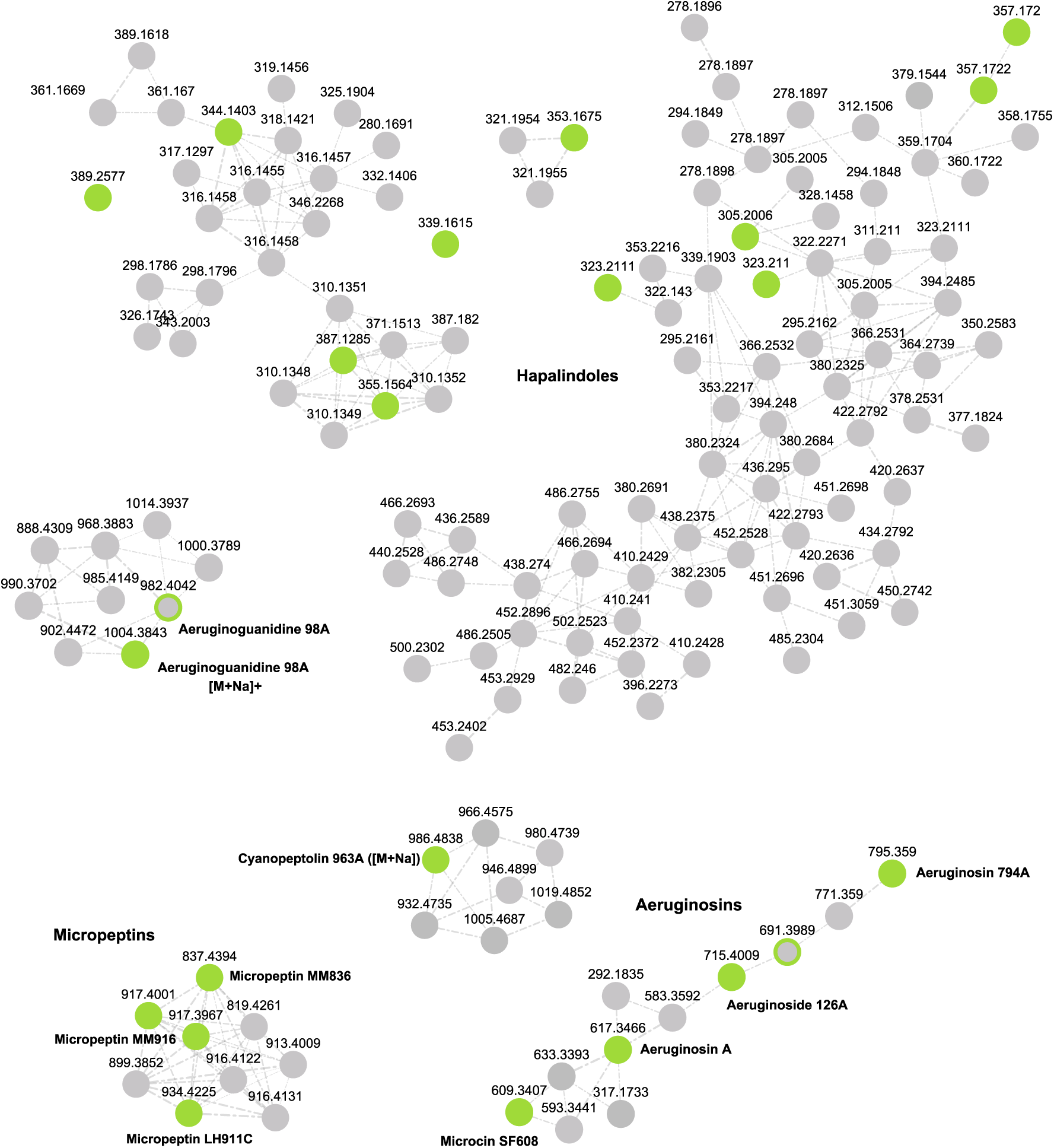
Feature Based Molecular Networking (FBMN) analysis of a sample set of cyanobacterial extracts with major compounds known a priori to the authors for chemical space visualization, pos. mode data. The compound annotation within this subset of the networking analysis was achieved through structure database search using the SIRIUS platform. Nodes connected for cosine scores ≥0.7, edge width correlating to cosine scores. Annotations derived from structure database searches are highlighted in green. Features without annotation, either corresponding to cyanobacterial SMs or uncharacterized compounds in general, are shown in gray. Shown clusters represent compound families lacking spectral reference data needed for confident annotation.

**Figure 7.**
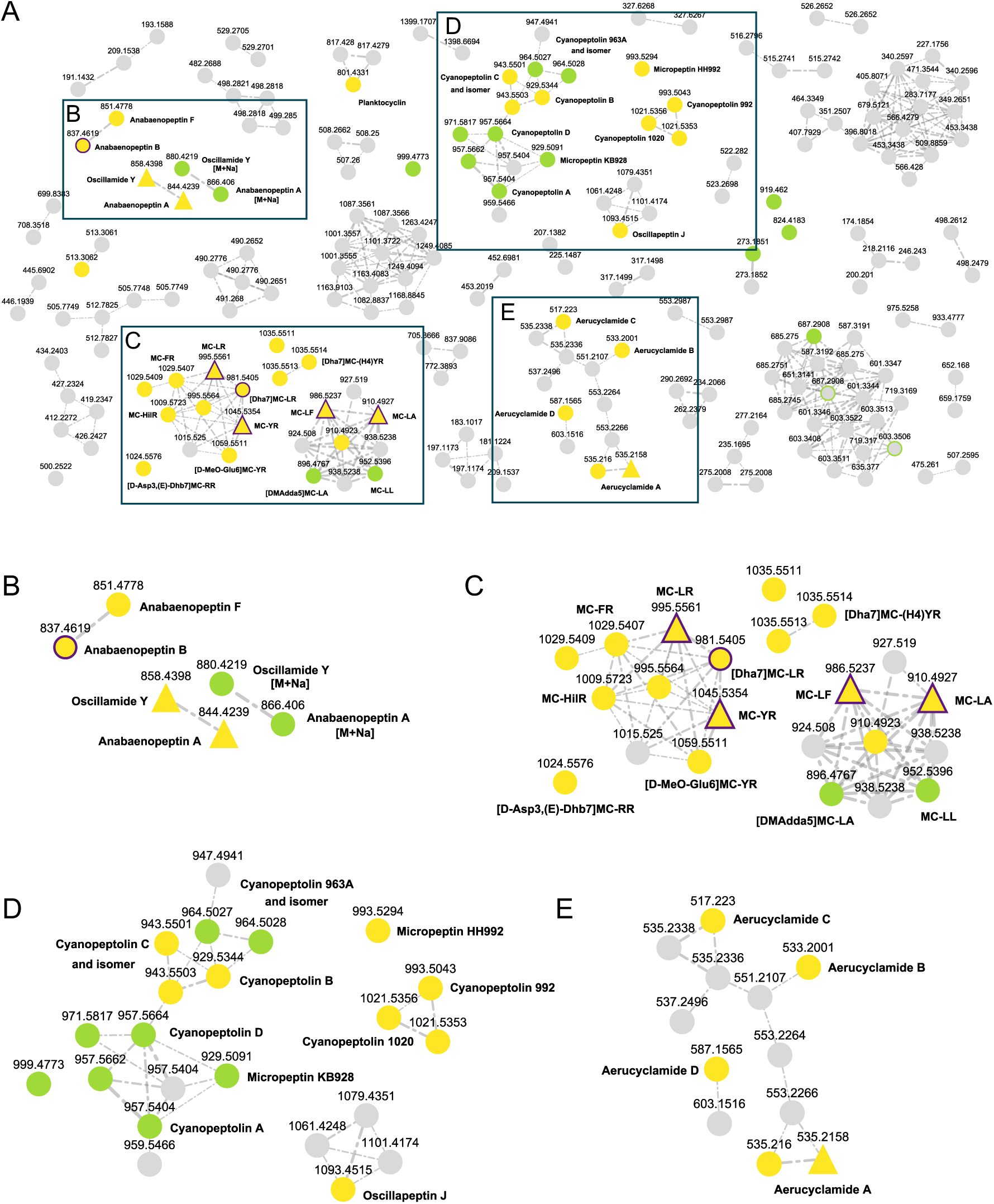
A. Feature Based Molecular Networking (FBMN) analysis (overview) of an unchracterized sample set of cyanobacterial extracts (Table 3) for chemical space visualization. Compound annotation was achieved through spectral library matching and structure database search using the SIRIUS platform. Nodes connected for cosine scores ≥0.6, edge width correlating to cosine scores. Clusters of the four main compound classes are highlighted. Annotations obtained through spectral library matching are highlighted in yellow, and those derived from structure database searches in green. Triangular nodes indicate spectra previously included in the MassBank database, while nodes outlined in purple represent features additionally annotated via GNPS. Features without annotation, either corresponding to cyanobacterial SMs or uncharacterized compounds in general, are shown in gray. **B** Cluster of anabaenopeptins. **C** Cluster of microcystins, displaying an increased spectral annotation rate compared to previous datasets. **D** Cluster of cyanopeptolins, for which spectral annotations were enabled by HRMS^2^ spectra recorded in this study. **E** Cluster of aerucyclamides, likewise demonstrating enhanced annotation coverage.

### Case study

As a case study, HPLC-HRMS data obtained from different cyanobacterial strains (Table S3) were used as an unbiased test set to assess which cyanobacterial SMs could be identified through spectral library matching without prior knowledge. Our aim was to evaluate whether the inclusion of new HRMS^2^ reference data could increase the confidence in the identification of known compounds and facilitate the annotation of previously uncharacterized chemical space of a dataset (Figure 7A). In total, FBMN analysis revealed 351 nodes, of which 300 were organized into clusters. Of these, 34 nodes were annotated via spectral library matching, while 16 were annotated through structure database searches. Notably, 25 of these 34 annotations could only be achieved when using the reference spectra generated in this study (Figure 7, Table S6). Two compounds were annotated with the newly acquired reference spectra as well as user-contributed spectra available on the GNPS platform. The inclusion of the new reference spectra led to a substantial increase in the overall annotation rate from roughly 2% to 10%. In conclusion, our newly acquired reference spectra were indispensable for correct annotation, as no other suitable data were available to support the identification of these compounds.

In both the proof-of-concept study and the case study, several structure database annotations were associated with features located in clusters already containing high-confidence spectral library matches, further supporting the reliability of these annotations. When cluster based annotation relationships are considered, like in the cyanopeptolin cluster (Figure 7D), the total annotation coverage across the dataset increased to approximately 13%. Moreover, clusters corresponding to compound classes such as anabaenopeptins, microcystins, and aerucyclamides could have already been identified based on existing reference data in public databases (Figure 7B, C, D). In contrast, the cyanopeptolin cluster and planktocyclin were only reliably annotated after incorporating the newly generated reference spectra. These high-confidence annotations can serve as anchors for the interpretation of additional nodes, allowing further compounds to be recognized based on exact mass matches or lower-confidence *in silico* predictions. This approach highlights how reliable reference spectra not only improve the confidence of individual annotations but also enable the systematic exploration of previously uncharacterized chemical space. High-confidence annotations derived from spectral library matching serve as key nodes within molecular networks. These key nodes increase the reliability of annotating connected features as via a structure database search (e.g., SIRIUS platform), or even when such features are supported only by exact mass matches (suspect lists), which represent a lower-confidence level of compound annotation, thereby enhancing dereplication.^50, 51^

A spectral-library similarity > 80 % was defined as a high-quality MS^2^ match. When this threshold was combined with the FBMN analysis - i.e., when the annotated compound was found within a cluster of structurally related SMs - a similarity > 70 % was also considered to provide good-confidence annotation. Scores ranging from 50 % to 70 % were regarded as “tentatively identified”, especially when the accurate mass measurement corroborated the proposed annotation. Features with a similarity < 50 % were excluded from further discussion, even though, as discussed above, the microginin exemplify a case where low thresholds might be linked to sub optimal acquisition parameters. This latter example, in particular, highlights that manual inspection and interpretation of the HRMS² data remain necessary; a careful, manual annotation of MS² spectra can provide stronger evidence for a confident identification.

### Conclusions and implications

The workflow to systematically record and process highly curated HRMS^2^ reference spectra for reposition in MassBank EU allowed us to increase available spectra for SMs from cyanobacteria from 11 to 150. This first attempt to generate reference spectra for metabolites in semi-purified extracts rather than only from reference materials required adjustments to the open-source scripts and a significant time commitment for manual quality control steps.

The new reference spectra are publicly available open access through MassBank EU and are automatically linked to other major spectral libraries, including the Human Metabolome Database (HMDB), MetaboLights, MassBank of North America (MoNA), and PubChem, the US EPA CompTox Dashboard, NORMAN Database System, RforMassSpectrometry. It is also used as a data source for searches in tools like GNPS and is included in the Metabolomics Spectrum Identifier Resolver. The reference spectra serve the purpose to obtain spectral matches in high-throughput non-target screening to identify tentative candidates and to improve dereplication efficiency. Compounds cannot be identified based on MS fragmentation alone, but the shorter list of matched tentative candidates can then be further investigated, e.g., by getting other reference samples from different sources to verify the spectra and retention times. Moreover, tentative candidates can serve as key nodes in molecular networking approaches such as GNPS and FBMN. This facilitates the systematic exploration of previously uncharacterized chemical space and supports accelerated dereplication strategies. The proof-of-concept and the case study revealed two key aspects: (i) several compounds were annotated through structure database searches but lacked corresponding experimental reference spectra, underscoring the need to further expand spectral databases; and (ii) in some cases, no annotation based on spectral library matching was obtained despite both the compound being present in the sample and a reference spectrum being available, highlighting that reference spectra may not always be perfectly representative or may not account for adduct formation influencing fragmentation behavior. Together, these findings emphasize the importance of continuously extending and refining the reference dataset in future work.

### Associated content

Supporting Information includes 6 Tables listing cyanobacterial SMs in MassBank EU deposited before 2025 (Table S1); strains used in the proof-of-concept study (Table S2); strains used in the case study (Table S3); 150 cyanobacterial SMs added to MassBank herein (Table S4); results of spectral library matching annotation of the proof-of-concept study (Table S5); and results of spectral library matching annotation of the case study (Table S6).

### Corresponding Author

*

### Author Contributions

**Franziska Schanbacher**: Conceptualisation, Analysis, Data processing, Writing Original Draft and Editing, Visualisations; **Anne Dax**: Analysis, Data processing, Writing Original Draft and Editing; **Michele Stravs**: Data processing, Editing Draft; **Timo Niedermeyer**: Conceptualisation, Editing Draft; **Elisabeth Janssen**: Conceptualisation, Writing Original Draft and Editing, Visualisations, Data Curation. All authors have given approval to the final version of the manuscript.

### Funding Sources

The mass spectrometers used in this study were partially funded by the German Research Foundation (DFG; INST 271/388-1, T.H.J.N.).

## Acknowledgements

We thank the CyanoMetDB collaborators for supplying reagents of metabolites with metadata and for performing final checks on deposited spectra: Daniel G. Beach, Dan Enke, Heike Enke, David P. Fewer, Fernanda Rios Jacinavicius, Jouni Jokela, Robert Konkel, Pedro Leão, Hanna Mazur-Marzec, Pearse McCarron, Antonella Miglione, Christopher O. Miles, Ernani Pinto, Marco Preto, Raphael Reher, Tania Shishido, Luciana Tartaglione, Mariana Torres, Matti Wahlsten.

