## Supplementary Materials for "Curation of mass spectrometry reference data for improved identification and dereplication of cyanobacterial specialized metabolites"

‡co-first authorship

### SUPPORTING INFORMATION

*The electronic supporting information contains 13 pages including 6 tables.*

**Table S1.** Cyanobacterial SMs in MassBank EU deposited before 2025 (last deposition 2020) with 14 unique compounds and 215 unique $HRMS^2$ spectra.

| compound name | MassBank EU assession numbers | precursor ion | resolution | HCDs (%) | deposition date | No spectra |
|---|---|---|---|---|---|---|
| **Microcystin-LR (MCLR)** | MSBNK-Eawag-EA299201 to MSBNK-Eawag-EA299217 | M+H | 7500 | 35, 15, 30, 45, 60, 75, 90 | 14.01.2014 | 7 |
| 29 spectra | MSBNK-Eawag-EA299208 to MSBNK-Eawag-EA299214 | M+H | 15000 | 15, 30, 45, 60,75, 90, 35 | 14.01.2014 | 7 |
| | MSBNK-Eawag-EQ299201 to MSBNK-Eawag-EQ299209 | M+H | 17500 | 15, 30,45, 60,75,90, 120, 150, 180 | 03.02.2020 | 9 |
| | MSBNK-Eawag-EQ299251 to MSBNK-Eawag-EQ299256 | M-H | 17500 | 15, 30, 45, 60, 75, 90 | 03.02.2020 | 6 |
| **Microcystin-LA (MCLA)** | MSBNK-Eawag-EQ324601 to MSBNK-Eawag-EQ324609 | M+H | 17500 | 15, 30,45, 60,75,90, 120, 150, 180 | 03.02.2020 | 9 |
| 15 spectra | MSBNK-Eawag-EQ324651 to MSBNK-Eawag-EQ324656 | M-H | 17500 | 15, 30, 45, 60, 75, 90 | 03.02.2020 | 6 |
| **Microcystin-LF (MCLF)** | MSBNK-Eawag-EQ324701 to MSBNK-Eawag-EQ324709 | M+H | 17500 | 15, 30,45, 60,75,90, 120, 150, 180 | 03.02.2020 | 9 |
| 18 spectra | MSBNK-Eawag-EQ324751 to MSBNK-Eawag-EQ324759 | M-H | 17500 | 15, 30,45, 60,75,90, 120, 150, 180 | 03.02.2020 | 9 |
| **Microcystin-LY (MCLY)** | MSBNK-Eawag-EQ324801 to MSBNK-Eawag-EQ324809 | M+H | 17500 | 15, 30,45, 60,75,90, 120, 150, 180 | 03.02.2020 | 9 |

| | | | | | | |
|---|---|---|---|---|---|---|
| 18 spectra | MSBNK-Eawag-EQ324851 to MSBNK-Eawag-EQ324859 | M-H | 17500 | 15, 30,45, 60,75,90, 120, 150, 180 | 03.02.2020 | 9 |
| **Microcystin-LW (MCLW)** | MSBNK-Eawag-EQ324901 to MSBNK-Eawag-EQ324909 | M+H | 17500 | 15, 30,45, 60,75,90, 120, 150, 180 | 03.02.2020 | 9 |
| 18 spectra | MSBNK-Eawag-EQ324951 to MSBNK-Eawag-EQ324959 | M-H | 17500 | 15, 30,45, 60,75,90, 120, 150, 180 | 03.02.2020 | 9 |
| **Microcystin-RR (MCRR)** | MSBNK-Eawag-EQ325001 to MSBNK-Eawag-EQ325006 | M+H | 35000 | 15, 30, 45, 60, 75, 90 | 25.08.2015 | 6 |
| 12 specrta | MSBNK-Eawag-EQ325051 to MSBNK-Eawag-EQ325054 | M-H | 17500 | 15, 30, 45, 60 | 03.02.2020 | 4 |
| | MSBNK-Eawag-EQ325056 | M-H | 35000 | 90 | 25.08.2015 | 1 |
| **Microcystin-YR (MCYR)** | MSBNK-Eawag-EQ325101 to MSBNK-Eawag-EQ325109 | M+H | 17500 | 15, 30,45, 60,75,90, 120, 150, 180 | 03.02.2020 | 9 |
| 18 spectra | MSBNK-Eawag-EQ325151 to MSBNK-Eawag-EQ325159 | M-H | 17500 | 15, 30,45, 60,75,90, 120, 150, 180 | 03.02.2020 | 9 |
| **[D-Asp3,E-Dhb7]-Microcystin-RR** | MSBNK-Eawag-EQ435801 to MSBNK-Eawag-EQ435809 | M+H | 17500 | 15, 30,45, 60,75,90, 120, 150, 180 | 03.02.2020 | 9 |
| 14 spectra | MSBNK-Eawag-EQ435851 to MSBNK-Eawag-EQ435855 | M-H | 17500 | 15, 30,45, 60,75 | 03.02.2020 | 5 |
| **Nodularin (Nodularin-R)** | MSBNK-Eawag-EQ325203 to MSBNK-Eawag-EQ325206 | M+H | 35000 | 45, 60, 75, 90 | 25.08.2015 | 4 |
| 10 spectra | MSBNK-Eawag-EQ325251 to MSBNK-Eawag-EQ325256 | M+H | 35000 | 15, 30, 45, 60, 75, 90 | 25.08.2015 | 6 |

| | | | | | | |
|---|---|---|---|---|---|---|
| **Anabaenopeptin A** | MSBNK-Eawag-EQ435601 to MSBNK-Eawag-EQ435609 | M+H | 17500 | 15, 30, 45, 60, 75, 90, 120, 150, 180 | 03.02.2020 | 9 |
| 15 spectra | MSBNK-Eawag-EQ435651 to MSBNK-Eawag-EQ435656 | M-H | 17500 | 15, 30, 45, 60, 75, 90 | 03.02.2020 | 6 |
| **Anabaenopeptin NZ857** | MSBNK-Eawag-EQ435901 to MSBNK-Eawag-EQ435909 | M+H | 17500 | 15, 30, 45, 60, 75, 90, 120, 150, 180 | 03.02.2020 | 9 |
| 15 spectra | MSBNK-Eawag-EQ435951 to MSBNK-Eawag-EQ435956 | M-H | 17500 | 15, 30, 45, 60, 75, 90 | 03.02.2020 | 6 |
| **Anabaenopeptin B** | MSBNK-Eawag-EQ436101 to MSBNK-Eawag-EQ436109 | M+H | 17500 | 15, 30, 45, 60, 75, 90, 120, 150, 180 | 03.02.2020 | 9 |
| 15 spectra | MSBNK-Eawag-EQ436151 to MSBNK-Eawag-EQ436156 | M-H | 17500 | 15, 30, 45, 60, 75, 90 | 03.02.2020 | 6 |
| **Oscillamide Y** | MSBNK-Eawag-EQ436001 to MSBNK-Eawag-EQ436009 | M+H | 17500 | 15, 30, 45, 60, 75, 90, 120, 150, 180 | 03.02.2020 | 9 |
| 9 spectra | | | | | | |
| **Aerucyclamide A** | MSBNK-Eawag-EQ436301 to MSBNK-Eawag-EQ436309 | M+H | 17500 | 15, 30, 45, 60, 75, 90, 120, 150, 180 | 03.02.2020 | 9 |
| 9 spectra | | | | | | |

**Table S2.** Strains used in the proof-of-concept study, with major compounds known a priori to the authors. Reference compounds were included to allow assessment of annotation and recovery rates in subsequent analyses within the proof-of-concept study.

| Strain-id | Genus | Expected SMs |
|---|---|---|
| id_1 | *Symphyonema* sp. | ambigols, tjipanazoles |
| id_2 | *Tolypothrix* sp. | cyanobacterins |
| id_3 | *Nostoc* sp. | nostotrebins |
| id_4 | *Nostoc* sp. | cryptophycins |
| id_5 | *Hapalosiphon* sp. | hapalindoles |
| id_6 | *Limnothrix* sp. | acutiphycin |
| id_7 | *Microcystis* sp. | aerucyclamides |
| id_8 | *Planktothrix* sp. | anabaenopeptins |
| id_9 | *Planktothrix* sp. | anabaenopeptins |
| id_10 | *Microcystis* sp. | microginins and microcystins |
| id_11 | *Cylindrospermum* sp. | cylindrofridins |
| id_12 | *Scytonema* sp. | scyptolins |
| id_13 | *Microcystis* sp. | microcystins |
| id_14 | *Nostoc* sp. | cryptophycins |
| id_15 | *Microcystis* sp. | microcystins |

**Table S3.** Strains used in the case study and their biomass extracts, with major compounds postulated by the authors based on manual $HRMS^2$ annotation.

| Strain-id | Genus (strain) | postulated SMs |
|---|---|---|
| id_16 | *Microcystis* sp. (PCC7806) | microcystins, cyanopeptolins, cyclamides |
| id_17 | *Planktothrix* sp. (*K-0576)* | microcystins, anabaenopeptin |
| id_18 | *Dolichospermum* sp. (*NIVA-CYA 269/6)* | microcystins, anabaenopeptin |
| id_19 | *Microcystis* sp. (UV006) | microcystins, cyanopeptolins, cyclamides |

**Table S4.** Cyanobacterial SMs represented with 150 unique compounds and 2911 unique HRMS$^2$ spectra (at 17500 resolution) in MassBank EU added herein and 11 compounds previously in Massbank (before 2025) represented with new spectra are marked with a hashtag (#), showing the number of spectra from M+2H, M+H and M-H precursor ions with scan mode "auto" and "40" (starting at *m/z* 40), as well as the total number of spectra deposited and the level of confidence for the compound identification.

| CyanoMetDB ID | Compound | Molecular Formula | Nr of spectra M+2H | Nr of M+H spectra; auto | Nr of M+H spectra; 40 | Nr of M-H spectra; auto | Nr of M-H spectra; 40 | Nr of total spectra | Level of confidence |
|---|---|---|---|---|---|---|---|---|---|
| 1823 # | MC-LR | C49H74N10O12 | 9 | 9 | 9 | 9 | 9 | 45 | 1 |
| 1802 # | MC-RR | C49H75N13O12 | 0 | 9 | 0 | 9 | 0 | 18 | 1 |
| 1844 # | MC-YR | C52H72N10O13 | 0 | 6 | 0 | 0 | 0 | 6 | 1 |
| 1894 # | MC-LA | C46H67N7O12 | 0 | 9 | 9 | 9 | 9 | 36 | 1 |
| 1916 # | MC-LY | C52H71N7O13 | 0 | 9 | 9 | 8 | 8 | 34 | 1 |
| 1962 # | [D-Asp3,(E)-Dhb7]MC-RR | C48H73N13O12 | 0 | 0 | 0 | 5 | n.a | 5 | 1 |
| 1953 | [D-Asp3]MC-LR | C48H72N10O12 | 9 | 9 | 0 | 9 | 0 | 27 | 1 |
| 1861 | [Dha7]MC-LR | C48H72N10O12 | 0 | 9 | 0 | 9 | 0 | 18 | 1 |
| 1879 | MC-HilR | C50H76N10O12 | 0 | 9 | 0 | 0 | 0 | 9 | 1 |
| 1972 | [D-Leu1]MC-LY | C55H77N7O13 | 0 | 9 | 0 | 9 | 0 | 18 | 1 |
| 2066 | MC-RY | C52H72N10O13 | 0 | 9 | 0 | 7 | 0 | 16 | 1 |
| 861 # | Nodularin-R | C41H60N8O10 | 0 | 9 | 9 | 9 | 9 | 36 | 1 |
| 867 # | Anabaenopeptin B | C41H60N10O9 | 9 | 9 | 0 | 9 | 0 | 27 | 1 |
| 1466 # | Aerucyclamide A | C24H34N6O4S2 | 0 | 7 | 0 | 0 | n.a | 7 | 1 |
| 1472 | Aerucyclamide B | C24H32N6O4S2 | 0 | 9 | 0 | 0 | n.a | 9 | 1 |
| 258 | Tychonamide A | C73H107N13O20 | 0 | 0 | 0 | 9 | n.a | 9 | 1 |
| 259 | Tychonamide B | C72H105N13O19 | 0 | 6 | 0 | 9 | n.a | 15 | 1 |
| 653 | Brunsvicamide B | C46H66N8O8 | 0 | 9 | 0 | 4 | n.a | 13 | 1 |
| 702 | Brunsvicamide C | C45H64N8O10 | 0 | 9 | 0 | 9 | n.a | 18 | 1 |
| 551 | Cyanopeptolin CP990 | C49H70N10O12 | 0 | 9 | 9 | 8 | 4 | 30 | 1 |
| 616 | Cyanopeptolin CP962 | C47H66N10O12 | 0 | 9 | 9 | 9 | 9 | 36 | 1 |
| 642 | Cyanopeptolin B | C46H72N8O12 | 0 | 18 | 9 | 4 | 0 | 31 | 1 |
| 983 | Carbamidocyclophane A | C38H54Cl4N2O8 | 0 | 0 | 0 | 4 | n.a | 4 | 1 |
| 978 | Carbamidocyclophane B | C38H55Cl3N2O8 | 0 | 0 | 0 | 4 | n.a | 4 | 1 |
| 972 | Carbamidocyclophane C | C38H56Cl2N2O8 | 0 | 0 | 0 | 4 | n.a | 4 | 1 |

| | | | | | | | | | |
|---|---|---|---|---|---|---|---|---|---|
| 969 | Carbamidocyclophane D | C38H57ClN2O8 | 0 | 0 | 0 | 5 | n.a | 5 | 1 |
| 963 | Carbamidocyclophane E | C38H58N2O8 | 0 | 0 | 0 | 6 | n.a | 6 | 1 |
| 214 | Molassamide | C48H66N8O13 | 0 | 7 | 0 | 7 | n.a | 14 | 1 |
| 2277 | Molassamide B | C48H65BrN8O13 | 0 | 8 | 0 | 6 | n.a | 14 | 1 |
| 2273 | Rivulariapeptolide 1185 | C61H87N9O15 | 0 | 9 | 0 | 9 | n.a | 18 | 1 |
| 2275 | Rivulariapeptolide 1121 | C56H83N9O15 | 0 | 9 | 0 | 9 | n.a | 18 | 1 |
| 2276 | Rivulariapeptolide 988 | C50H68N8O13 | 0 | 9 | 0 | 3 | n.a | 12 | 1 |
| 2364 | Cylindrocyclophane B | C38H58O7 | 0 | 0 | 0 | 5 | n.a | 5 | 1 |
| 2365 | Cylindrocyclophane D | C40H60O8 | 0 | 0 | 0 | 6 | n.a | 6 | 1 |
| 2340 | Insulapeptolide D | C48H75N9O12 | 0 | 9 | 0 | 5 | n.a | 14 | 1 |
| 2341 | Insulapeptolide E | C51H72N8O14 | 0 | 9 | 0 | 9 | n.a | 18 | 1 |
| 2342 | Insulapeptolide F | C50H70N8O14 | 0 | 9 | 0 | 8 | n.a | 17 | 1 |
| 2343 | Insulapeptolide G | C50H70N8O13 | 0 | 9 | 0 | 7 | n.a | 16 | 1 |
| 2344 | Insulapeptolide H | C51H72N8O13 | 0 | 9 | 0 | 7 | n.a | 16 | 1 |
| 21 | Tutuilamide A | C51H69ClN8O12 | 0 | 9 | 0 | 8 | n.a | 17 | 1 |
| 155 | Balticidin C | C75H121N11O36 | 0 | 9 | 0 | 0 | n.a | 9 | 1 |
| 1213 | Gallinamide A | C31H52N4O7 | 0 | 9 | 0 | 0 | n.a | 9 | 1 |
| 1240 | Aeruginosin NOL3 | C30H48N6O6 | 0 | 9 | 9 | 3 | 4 | 25 | 1 |
| 1339 | Acutiphycin | C27H44O7 | 0 | 0 | 0 | 8 | n.a | 8 | 1 |
| 525 | Nostotrebin 6 | C50H38O10 | 0 | 7 | 0 | 0 | n.a | 7 | 1 |
| 923 | Nostocyclopeptide A2 | C40H54N8O9 | 0 | 9 | 9 | 9 | 4 | 31 | 1 |
| 924 | Kasumigamide | C40H50N8O9 | 0 | 8 | 0 | 6 | n.a | 14 | 1 |
| 2367 | Nostolactone 4 | C25H20O6 | 0 | 9 | 0 | 8 | n.a | 17 | 1 |
| 2520 | Anhydrocyanobacterin | C23H21ClO5 | 0 | 9 | 0 | 0 | n.a | 9 | 1 |
| 760 # | Anabaenopeptin A | C44H57N7O10 | 0 | 8 | 0 | 7 | n.a | 15 | 2a |
| 713 # | Oscillamide Y | C45H59N7O10 | 0 | 5 | 0 | 4 | n.a | 9 | 2a |
| 859 | Anabaenopeptin C | C41H60N8O9 | 0 | 8 | 0 | 7 | n.a | 15 | 2a |
| 1118 | Bartoloside E | C34H58Cl2O6 | 0 | 9 | 9 | 7 | 0 | 25 | 2a |
| 993 | Bartoloside F | C37H64Cl2O6 | 0 | 9 | 9 | 7 | 4 | 29 | 2a |
| 1027 | Bartoloside G | C36H63ClO6 | 0 | 9 | 9 | 7 | 0 | 25 | 2a |
| 1031 | Bartoloside I | C36H61Cl3O6 | 0 | 9 | 9 | 7 | 0 | 25 | 2a |
| 1117 | Bartoloside J | C34H59ClO6 | 0 | 9 | 9 | 7 | 0 | 25 | 2a |
| 1079 | Bartoloside K | C35H60Cl2O6 | 0 | 9 | 9 | 0 | 0 | 18 | 2a |

| | | | | | | | | | |
|---|---|---|---|---|---|---|---|---|---|
| 352 | Lyngbyazothrin A | C62H96N12O19 | 0 | 9 | 0 | 6 | n.a | 15 | 2a |
| 355 | Lyngbyazothrin B | C61H94N12O18 | 0 | 9 | 0 | 6 | n.a | 15 | 2a |
| 326 | Lyngbyazothrin D | C73H107N13O21 | 0 | 9 | 0 | 2 | n.a | 11 | 2a |
| 248 | Tjipanazol D | C18H10Cl2N2 | 0 | 0 | 0 | 9 | n.a | 9 | 2a |
| 255 | Tjipanazol I | C18H11ClN2 | 0 | 0 | 0 | 9 | n.a | 9 | 2a |
| 2518 | Tjipanazol L | C20H9Cl2N3O2 | 0 | 0 | 0 | 9 | n.a | 9 | 2a |
| 2519 | Tjipanazol M | C20H10ClN3O2 | 0 | 0 | 0 | 8 | n.a | 8 | 2a |
| 41 | Portoamide A | C74H109N13O22 | 0 | 9 | 9 | 0 | 0 | 18 | 2a |
| 42 | Portoamide B | C73H107N13O21 | 0 | 9 | 9 | 0 | 0 | 18 | 2a |
| 173 | Portoamide C | C62H96N12O19 | 0 | 8 | 9 | 9 | 9 | 35 | 2a |
| 174 | Portoamide D | C61H94N12O18 | 0 | 8 | 9 | 9 | 9 | 35 | 2a |
| 684 | Scyptolin A | C45H69ClN8O14 | 0 | 9 | 0 | 3 | n.a | 12 | 2a |
| 464 | Scyptolin B | C52H80ClN9O16 | 0 | 7 | 0 | 3 | n.a | 10 | 2a |
| 190 | Scytocyclamide B | C63H110N14O19 | 0 | 9 | 0 | 6 | n.a | 15 | 2a |
| 191 | Scytocyclamide C | C63H110N14O18 | 0 | 9 | 0 | 2 | n.a | 11 | 2a |
| 1104 | Cryptophycin A | C35H43ClN2O8 | 0 | 9 | 0 | 6 | n.a | 15 | 2a |
| 1469 | Aerucyclamide C | C24H32N6O5S | 0 | 18 | 9 | 0 | 0 | 27 | 2a |
| 149 | Sphaerocyclamide | C46H63N9O11 | 0 | 9 | 9 | 9 | 7 | 34 | 2a |
| 15 | Nocuolin A | C16H30N2O3 | 0 | 9 | 9 | 9 | 9 | 36 | 2a |
| 900 | Cylindrofridin C | C40H61ClO8 | 0 | 0 | 0 | 5 | n.a | 5 | 2a |
| 904 | Cyanostatin B | C40H59N5O9 | 0 | 7 | 0 | 7 | n.a | 14 | 2a |
| 961 | Cylindrofridin B | C38H59ClO7 | 0 | 0 | 0 | 5 | n.a | 5 | 2a |
| 1282 | Namalide B | C29H45N5O7 | 0 | 9 | 9 | 9 | 9 | 36 | 2a |
| 811 | Schizopeptin 791 | C42H61N7O8 | 0 | 9 | 9 | 6 | 4 | 28 | 2a |
| 1481 | Hierridin B | C23H40O3 | 0 | 9 | 8 | 9 | 9 | 35 | 2a |
| 1694 | 7-Deoxy-Cylindrospermopsin | C15H21N5O6S | 0 | 9 | 0 | 9 | 0 | 18 | 2a |
| 2515 | Monomethylaetokthonostatin | C42H73N5O7 | 0 | 9 | 0 | 0 | n.a | 9 | 2a |
| 140 | Pseudospumigin A | C31H44N6O7 | 0 | 9 | 9 | 0 | 0 | 18 | 2a |
| 1800 | MC-FR | C52H72N10O12 | 0 | 7 | 0 | 6 | 0 | 13 | 2b |
| 1835 | MC-FL | C52H71N7O12 | 0 | 9 | 9 | 7 | 8 | 33 | 2b |
| 1869 | MC-YL | C52H71N7O13 | 0 | 9 | 0 | 7 | 0 | 16 | 2b |
| 1877 | [D-MeO-Glu6]MC-LR | C50H76N10O12 | 0 | 9 | 9 | 8 | 7 | 33 | 2b |
| 1895 | [D-Asp3]MC-RY | C51H70N10O13 | 0 | 9 | 9 | 9 | 9 | 36 | 2b |

| | | | | | | | | | |
|---|---|---|---|---|---|---|---|---|---|
| 1950 | [D-Asp3,Dha7]MC-LR | C47H70N10O12 | 0 | 9 | 0 | 5 | n.a | 14 | 2b |
| 1997 | [D-Leu1]MC-LR | C52H80N10O12 | 0 | 9 | 9 | 8 | 6 | 32 | 2b |
| 2036 | [D-Met(O)1]MC-LR | C51H78N10O13S | 0 | 5 | 0 | 0 | 0 | 5 | 2b |
| 668 | Anabaenopeptin 871 | C46H61N7O10 | 0 | 5 | 0 | 6 | 0 | 11 | 2b |
| 805 | Anabaenopeptin F | C42H62N10O9 | 0 | 9 | 9 | 9 | 9 | 36 | 2b |
| 809 | Anabaenopeptin 807 | C42H61N7O9 | 0 | 0 | 0 | 4 | 0 | 4 | 2b |
| 467 | Cyanopeptolin CP1048 | C52H76N10O13 | 0 | 9 | 9 | 9 | 9 | 36 | 2b |
| 512 | Cyanopeptolin 1020 | C50H72N10O13 | 0 | 9 | 9 | 9 | 9 | 36 | 2b |
| 532 | Cyanopeptolin 1014 | C49H78N10O13 | 0 | 9 | 9 | 4 | 4 | 26 | 2b |
| 555 | Cyanopeptolin 963A | C49H69N7O13 | 0 | 0 | 0 | 9 | 0 | 9 | 2b |
| 603 | Cyanopeptolin C | C47H74N8O12 | 0 | 15 | 0 | 13 | 0 | 28 | 2b |
| 605 | Micropeptin K139 | C47H74N10O13 | 0 | 9 | 9 | 9 | 9 | 36 | 2b |
| 613 | Oscillapeptin J | C47H68N10O18S | 0 | 9 | 9 | 5 | 9 | 32 | 2b |
| 638 | Cyanopeptolin 972 | C46H72N10O13 | 0 | 9 | 9 | 9 | 9 | 36 | 2b |
| 699 | Nodulapeptin 879 | C45H65N7O11 | 0 | 5 | 0 | 3 | 0 | 8 | 2b |
| 733 | [Met6] Nodulapeptin C | C44H65N7O9S2 | 0 | 9 | 9 | 6 | 5 | 29 | 2b |
| 735 | Nodulapeptin 883a | C44H65N7O8S2 | 0 | 4 | 0 | 4 | 0 | 8 | 2b |
| 748 | Nodulapeptin B | C44H63N7O12S | 0 | 9 | 0 | 6 | 0 | 15 | 2b |
| 750 | Nodulapeptin C | C44H63N7O11S | 0 | 9 | 9 | 9 | 8 | 35 | 2b |
| 752 | Nodulapeptin 881a | C44H63N7O10S | 0 | 9 | 7 | 6 | 4 | 26 | 2b |
| 2240 | Microginin 299A | C45H67ClN6O10 | 0 | 9 | 9 | 9 | 9 | 36 | 2b |
| 2241 | Microginin 299B | C45H66Cl2N6O10 | 0 | 9 | 9 | 5 | 5 | 28 | 2b |
| 2242 | Microginin 299C | C45H68N6O10 | 0 | 9 | 9 | 9 | 9 | 36 | 2b |
| 975 | Microginin FR5 | C38H55N5O9 | 0 | 7 | 0 | 0 | 0 | 7 | 2b |
| 1019 | Microginin 761B | C37H52ClN5O10 | 0 | 9 | 0 | 4 | 0 | 13 | 2b |
| 948 | Microginin 757 | C39H59N5O10 | 0 | 9 | 0 | 0 | 0 | 9 | 2b |
| 847 | Microginin 770 | C41H63N5O9 | 0 | 9 | 9 | 9 | 9 | 36 | 2b |
| 897 | Microginin SD755 | C40H61N5O9 | 0 | 9 | 0 | 0 | 0 | 9 | 2b |
| 898 | Microginin 756 | C40H61N5O9 | 0 | 9 | 9 | 9 | 9 | 36 | 2b |
| 947 | Nostoginin BN741 | C39H59N5O9 | 0 | 9 | 9 | 0 | 9 | 27 | 2b |
| 1221 | Spumigin A | C31H44N6O7 | 0 | 9 | 9 | 3 | 3 | 24 | 2b |
| 1250 | Spumigin D | C30H42N6O7 | 0 | 9 | 9 | 0 | 0 | 18 | 2b |
| 1256 | Spumigin F | C30H40N6O7 | 0 | 9 | 9 | 0 | 0 | 18 | 2b |

| | | | | | | | | | |
|---|---|---|---|---|---|---|---|---|---|
| 1231 | Spumigin G | C31H42N6O6 | 0 | 9 | 9 | 0 | 0 | 18 | 2b |
| 1504 | Nostosin A | C22H35N5O5 | 0 | 9 | 9 | 0 | 0 | 18 | 2b |
| 1503 | Nostosin B | C22H37N5O5 | 0 | 9 | 9 | 0 | 0 | 18 | 2b |
| 2163 | Nostocyclopeptide Ncp-E1-L | C39H54N8O10 | 0 | 9 | 9 | 9 | 9 | 36 | 2b |
| 2165 | Nostocyclopeptide Ncp-E2-L | C36H56N8O10 | 0 | 9 | 9 | 9 | 9 | 36 | 2b |
| 1245 | Aeruginosin NAL2 | C30H46N6O6 | 0 | 9 | 9 | 6 | 6 | 30 | 2b |
| 1275 | Oscillaginin A | C29H47ClO8N4 | 0 | 9 | 0 | 7 | 0 | 16 | 2b |
| 1332 | Muscoride A | C28H40N4O5 | 0 | 9 | 9 | 0 | 0 | 18 | 2b |
| 1402 | Aerucyclamide D | C26H30N6O4S3 | 0 | 9 | 9 | 3 | 0 | 21 | 2b |
| 137 | Namalide D | C29H45N5O6 | 0 | 9 | 9 | 8 | 8 | 34 | 2b |
| 260 | Muscoride B | C31H41N5O6 | 0 | 9 | 9 | 0 | 0 | 18 | 2b |
| 266 | Anabaenolysin A | C28H38N4O8 | 0 | 9 | 2 | 9 | 8 | 28 | 2b |
| 421 | Planktopeptin BL 1125 | C54H79N9O17 | 0 | 9 | 0 | 9 | 0 | 18 | 2b |
| 939 | Planktocyclin | C39H60N8O8S | 0 | 9 | 0 | 0 | 0 | 9 | 2b |
| 584 | Cyanopeptolin 992 | C48H68N10O13 | 0 | 9 | 9 | 4 | 0 | 22 | 3 |
| 912 | Microginin GH787 | C40H58ClN5O9 | 0 | 9 | 9 | 8 | 0 | 26 | 3 |
| **Isomer Group_C24H34N6O6S** | | | C24H34N6O6S | 0 | 0 | 0 | 9 | 0 | 9 |
| 1471 | Microcyclamide 7806A | C24H34N6O6S | | | | | | | |
| 1465 | Microcyclamide 7806B | C24H34N6O6S | | | | | | | |
| **Isomer Group_C42H61N7O10S** | | | C42H61N7O10S | 0 | 9 | 0 | 6 | 0 | 15 |
| 813 | Nodulapeptin 855a | C42H61N7O10S | | | | | | | |
| 814 | Nodulapeptin 855b | C42H61N7O10S | | | | | | | |
| **Isomer Group_C41H61N5O9** | | | C41H61N5O9 | 0 | 9 | 9 | 4 | 0 | 22 |
| 857 | Microginin 767 | C41H61N5O9 | | | | | | | |
| 858 | Microginin KR767 | C41H61N5O9 | | | | | | | |
| **Isomer Group_C31H44N6O7** | | | C31H44N6O7 | 0 | 9 | 9 | 0 | 0 | 18 |
| 1222 | Dihydrospumigin K | C31H44N6O7 | | | | | | | |
| 1223 | Dihydrospumigin L | C31H44N6O7 | | | | | | | |
| **Isomer Group_C31H42N6O7** | | | C31H42N6O7 | 0 | 9 | 9 | 4 | 3 | 25 |
| 1228 | Spumigin E | C31H42N6O7 | | | | | | | |
| 1229 | Spumigin K | C31H42N6O7 | | | | | | | |
| 1230 | Spumigin L | C31H42N6O7 | | | | | | | |

| | | | | | | | | | |
|---|---|---|---|---|---|---|---|---|---|
| **Isomer Group_C31H42N6O7** | | | C30H42N6O7 | 0 | 9 | 9 | 0 | 0 | 18 |
| 1251 | Dihydrospumigin M | C30H42N6O7 | | | | | | | |
| 1252 | Dihydrospumigin N | C30H42N6O7 | | | | | | | |
| **Isomer Group_C51H78N10O12** | | | C51H78N10O12 | 0 | 0 | 0 | 5 | 5 | 10 |
| 1834 | [Leu1,D-Asp3]MC-LR | C51H78N10O12 | | | | | | | |
| 1911 | [Leu1,Dha7]MC-LR | C51H78N10O12 | | | | | | | |
| **Isomer Group_C51H78N10O12** | | | C51H74N10O13 | 0 | 7 | 0 | 0 | n.a | 7 |
| 1947 | [Dha7]MC-(H4)YR | C51H74N10O13 | | | | | | | |
| 1948 | [D-Asp3]MC-(H4)YR | C51H74N10O13 | | | | | | | |
| 1949 | [DMAdda5]MC-(H4)YR | C51H74N10O13 | | | | | | | |
| **Isomer Group_C53H74N10O13** | | | C53H74N10O13 | 0 | 7 | 6 | 8 | 7 | 28 |
| 2014 | [D-Glu(OMe)6]MC-YR | C53H74N10O13 | | | | | | | |
| 2015 | MCHtyR | C53H74N10O13 | | | | | | | |
| 2016 | [D-Asp3,D-Glu(OMe)6]MC-HtyR | C53H74N10O13 | | | | | | | |
| **Isomer Group_C50H72N8O13** | | | C50H72N8O13 | 0 | 7 | 0 | 0 | 0 | 7 |
| 509 | Micropeptin HH992 | C50H72N8O13 | | | | | | | |
| 510 | Micropeptin KB992 | C50H72N8O13 | | | | | | | |
| 511 | Loggerpeptin A | C50H72N8O13 | | | | | | | |

**Table S5.** Spectral library matching annotation results obtained for the biomass extracts analyzed in the proof-of-concept study (Table S2). Mass deviation from exact mass in parentheses. Annotations exclusively based on newly recorded reference spectra are highlighted (*).

| specialized metabolite (expected) | specialized metabolite (annotated) | *m/z* (accurate mass) | *m/z* (exact mass) | mol. formula | spectral library matching |
|---|---|---|---|---|---|
| ambigols, tjipanazoles | Tjipanazole D* | 323.0145 [M-H]$^-$ | 323.0148 [M-H]$^-$ | $C_{18}H_{10}C_{12}N_2$ (Δ 0.9 ppm) | 78% |
| | Tjipanazole I* | 289.0539 [M-H]$^-$ | 289.0538 [M-H]$^-$ | $C_{18}H_{11}ClN_2$ (Δ 0.3 ppm) | 77% |
| Cyanobacterin and analogues | Anhyrocyanobactin (isomers 1 + 2)* | 413.1149 [M+H]$^+$ | 413.1150 [M+H]$^+$ | $C_{23}H_{21}ClO_5$ (Δ 0.2 ppm) | 81% / 82% |
| Nostotrebin 6 and related compounds | Nostolacton 4* | 417.1329 [M+H]$^+$ | 417.1333 [M+H]$^+$ | $C_{25}H_{20}O_6$ (Δ 0.2 ppm) | 87% |
| | Nostotrebin 6* | 799.2533 [M+H]$^+$ | 799.2538 [M+H]$^+$ | $C_{50}H_{38}O_{10}$ (Δ 0.6 ppm) | 68% |
| | (3 isomers/analogues) | 799.2505 [M+H]$^+$ | 799.2538 [M+H]$^+$ | $C_{50}H_{38}O_{10}$ (Δ 4.1 ppm) | 66% |
| | | 799.2531 [M+H]$^+$ | 799.2538 [M+H]$^+$ | $C_{50}H_{38}O_{10}$ (Δ 0.9 ppm) | 88% |
| Cryptophycins | Cryptophycin A* | 655.2775 [M+H]$^+$ | 655.2781 [M+H]$^+$ | $C_{35}H_{43}ClN_2O_8$ (Δ 0.9 ppm) | 88% |
| Acutiphycin | no annotation | - | - | - | - |
| Aerucyclamides | Aerucyclamide A | 535.2156 [M+H]$^+$ | 535,2156 [M+H]$^+$ | $C_{24}H_{34}N_6O_4S_2$ (Δ 0.0 ppm) | 82% |
| | (3 isomers) | 535.2174 [M+H]$^+$ | 535,2156 [M+H]$^+$ | $C_{24}H_{34}N_6O_4S_2$ (Δ 3.4 ppm) | 83% |
| | | 535.2161 [M+H]$^+$ | 535,2156 [M+H]$^+$ | $C_{24}H_{34}N_6O_4S_2$ (Δ 0.9 ppm) | 79% |
| | Aerucyclamide C* | 517.2227 [M+H]$^+$ | 517.2228 [M+H]$^+$ | $C_{24}H_{32}N_6O_5S$ (Δ 0.2 ppm) | 83% |
| | (2 isomers) | 517.2224 [M+H]$^+$ | 517.2228 [M+H]$^+$ | $C_{24}H_{32}N_6O_5S$ (Δ 0.8 ppm) | 85% |
| Anabaenopeptins | Anabaenopeptin A | 844.4241 [M+H]$^+$ | 844,4240 [M+H]$^+$ | $C_{44}H_{57}N_7O_{10}$ (Δ 0.2 ppm) | 94% |
| | Anabaenopeptin C* | 809.4542 [M+H]$^+$ | 809.4556 [M+H]$^+$ | $C_{41}H_{60}N_8O_9$ (Δ 1.7 ppm) | 85% |
| | Anabaenopeptin B | 837.4615 [M+H]$^+$ | 837.4617 [M+H]$^+$ | $C_{41}H_{60}N_{10}O_9$ (Δ 0.3 ppm) | 94% |
| | Anabaenopeptin F* | 851.4772 [M+H]$^+$ | 851,4774 [M+H]$^+$ | $C_{42}H_{62}N_{10}O_9$ (Δ 0.2 ppm) | 92% |
| | Oscillamide Y | 858.4399 [M+H]$^+$ | 858.4396 [M+H]$^+$ | $C_{45}H_{59}N_7O_{10}$ (Δ 0.3 ppm) | 79% |
| Microginins | Microginin SD755* | 756.4548 [M+H]$^+$ | 756.4542 [M+H]$^+$ | $C_{40}H_{61}N_5O_9$ (Δ 0.8 ppm) | 32% |
| | Microginin 742A* | 742.4383 [M+H]$^+$ | 742.4386 [M+H]$^+$ | $C_{39}H_{59}N_5O_9$ (Δ 0.4 ppm) | 24% |
| Cylindrofridins | Cylindrofridin B* | 661.3861 [M-H]$^-$ | 661.3876 [M-H]$^-$ | $C_{38}H_{59}ClO_7$ (Δ 2.3 ppm) | 68% |
| | Cylindrofridin C* | 703.3989 [M-H]$^-$ | 703.3982 [M-H]$^-$ | $C_{40}H_{61}ClO_8$ (Δ 1.0 ppm) | 76% |
| Scyptolin A, B | Scyptolin A* | 979.4552 [M-H]$^-$ | 979.4549 [M-H]$^-$ | $C_{45}H_{69}ClN_8O_{14}$ (Δ 0.3 ppm) | 61% |
| | Scyptolin B* | 1120.5383 M-H]$^-$ | 1120.5339 M-H]$^-$ | $C_{52}H_{80}ClN_9O_{16}$ (Δ 3.9 ppm) | 56% |
| Microcystins | MC-LR | 995.5557 [M+H]$^+$ | 995.5560 [M+H]$^+$ | $C_{49}H_{74}N_{10}O_{12}$ (Δ 0.3 ppm) | 83% |
| | MC-YR | 1045.5353 [M+H]$^+$ | 1045.5353 [M+H]$^+$ | $C_{52}H_{72}N_{10}O_{13}$ (Δ 0.0 ppm) | 80% |
| | [D-Asp3]MC-LR | 981.5403 [M+H]$^+$ | 981.5404 [M+H]$^+$ | $C_{48}H_{72}N_{10}O_{12}$ (Δ 0.1 ppm) | 72% |
| | MC-LA | 910.4916 [M+H]$^+$ | 910.4920 [M+H]$^+$ | $C_{46}H_{67}N_7O_{12}$ (Δ 0.4 ppm) | 87% |
| | MC-LW | 1025.5348 [M+H]$^+$ | 1025.5342 [M+H]$^+$ | $C_{54}H_{70}N_8O_{12}$ (Δ 0.6 ppm) | 68% |
| SMs not known a priori | Cylindrocyclophane D* | 667.4236 [M-H]$^-$ | 667.4215 [M-H]$^-$ | $C_{40}H_{60}O_8$ (Δ 3.1 ppm) | 77% |
| | Planktocyclin * | 801.4325 [M+H]$^+$ | 801,4327 [M+H]$^+$ | $C_{39}H_{60}N_8O_8S$ (Δ 0.2 ppm) | 62% |

**Table S6.** Spectral library matching annotation results obtained for the biomass extracts analyzed in the case study (Table S3). Mass deviation from exact mass in parentheses.

| specialized metabolite (annotated) | *m/z* [M+H]$^+$ (accurate mass) | *m/z* [M+H]$^+$ (exact mass) | mol. formula | spectral library matching |
|---|---|---|---|---|
| Aerucyclamide A | 535.2158 | 535.156 | $C_{24}H_{34}N_6O_4S_2$ (Δ 0.4 ppm) | 85% |
| (2 isomers) | 535.2160 | 535.2156 | $C_{24}H_{34}N_6O_4S_2$ (Δ 0.7 ppm) | 85% |
| Aerucyclamide B | 533.2001 | 533.1999 | $C_{24}H_{32}N_6O_4S_2$ (Δ 0.4 ppm) | 90% |
| Aerucyclamide C | 517.2230 | 517.2228 | $C_{24}H_{32}N_6O_5S$ (Δ 0.4 ppm) | 95% |
| Aerucyclamide D | 587.1565 | 587.1563 | $C_{26}H_{30}N_6O_4S_3$ (Δ 0.3 ppm) | 85% |
| Anabaenopeptin A | 844.4239 | 844.4240 | $C_{44}H_{57}N_7O_{10}$ (Δ 0.1 ppm) | 84% |
| Anabaenopeptin B | 837.4619 | 837.4617 | $C_{41}H_{60}N_{10}O_9$ (Δ 0.2 ppm) | 98% |
| Anabaenopeptin F | 851.4778 | 851.4774 | $C_{42}H_{62}N_{10}O_9$ (Δ 0.5 ppm) | 96% |
| Oscillamide Y | 858.4398 | 858.4396 | $C_{45}H_{59}N_7O_{10}$ (Δ 0.2 ppm) | 74% |
| Oscillapeptin J | 1093.4515 | 1093.4507 | $C_{47}H_{68}N_{10}O_{18}S$ (Δ 0.7 ppm) | 97% |
| Cyanopeptolin 1020 | 1021.5353 | 1021.5358 | $C_{50}H_{72}N_{10}O_{13}$ (Δ 0.0 ppm) | 86% |
| (2 isomers) | 1021.5356 | 1021.5358 | $C_{50}H_{72}N_{10}O_{13}$ (Δ 0.2 ppm) | 84% |
| Cyanopeptolin 992 | 993.5043 | 993.5040 | $C_{48}H_{68}N_{10}O_{13}$ (Δ 0.3 ppm) | 84% |
| Cyanopeptolin B | 929.5340 | 929.5342 | $C_{46}H_{72}N_8O_{12}$ (Δ 0.2 ppm) | 94% |
| Cyanopeptolin C | 943.5503 | 943.5499 | $C_{47}H_{74}N_8O_{12}$ (Δ 0.4 ppm) | 92% |
| (2 isomers) | 943.5501 | 943.5499 | $C_{47}H_{74}N_8O_{12}$ (Δ 0.2 ppm) | 81% |
| Micropeptin HH992 | 993.5294 | 993.5292 | $C_{50}H_{72}N_8O_{13}$ (Δ 0.2 ppm) | 95% |
| MC-LR | 995.5561 | 995.5560 | $C_{49}H_{74}N_{10}O_{12}$ (Δ 0.1 ppm) | 89% |
| MC-YR | 1045.5354 | 1045.5353 | $C_{52}H_{72}N_{10}O_{13}$ (Δ 0.1 ppm) | 73% |
| MC-FR* | 1029.5407 | 1029.5404 | $C_{52}H_{72}N_{10}O_{12}$ (Δ 0.3 ppm) | 76% |
| [Dha7]MC-LR | 981.5405 | 981.5404 | $C_{48}H_{72}N_{10}O_{12}$ (Δ 0.1 ppm) | 93% |
| [D-MeO-Glu6]MC-YR | 1059.5511 | 1059.5511 | $C_{53}H_{74}N_{10}O_{13}$ (Δ 0.1 ppm) | 87% |
| [D-Asp3,(E)-Dhb7]MC-RR | 1024.5576 | 1024.5574 | $C_{48}H_{73}N_{13}O_{12}$ (Δ 0.2 ppm) | 77% |
| MC-HilR | 1009.5723 | 1009.5717 | $C_{50}H_{76}N_{10}O_{12}$ (Δ 0.6 ppm) | 89% |
| MC-LA | 910.4927 | 910.4920 | $C_{46}H_{67}N_7O_{12}$ (Δ 0.8 ppm) | 94% |
| MC-LF | 986.5237 | 986.5233 | $C_{52}H_{71}N_7O_{12}$ (Δ 0.4 ppm) | 92% |
| Planktocyclin | 801.4331 | 801.4327 | $C_{39}H_{60}N_8O_8S$ (Δ 0.5 ppm) | 83% |
